# Strong selective environments determine evolutionary outcome in time-dependent fitness seascapes

**DOI:** 10.1101/2020.12.01.406181

**Authors:** Johannes Cairns, Florian Borse, Tommi Mononen, Teppo Hiltunen, Ville Mustonen

## Abstract

The impact of fitness landscape features on evolutionary outcomes has attracted considerable interest in recent decades. However, evolution often occurs under time-dependent selection in so-called fitness seascapes where the landscape is under flux. Fitness seascapes are an inherent feature of natural environments, where the landscape changes owing both to the intrinsic fitness consequences of previous adaptations and extrinsic changes in selected traits caused by new environments. The complexity of such seascapes may curb the predictability of evolution. However, empirical efforts to test this question utilising a comprehensive set of regimes are lacking. Here we employed an *in vitro* microbial model system to investigate differences in evolutionary outcomes between time-invariant and -dependent environments, including all possible temporal permutations, with three subinhibitory antimicrobials and a viral parasite (phage) as selective agents. Expectedly, time-invariant environments caused stronger directional selection for resistances compared to time-dependent environments. Intriguingly, however, multidrug resistance outcomes in both cases were largely driven by two strong selective agents (rifampicin and phage) out of four agents in total. These agents either caused cross-resistance or obscured the phenotypic effect of other resistance mutations, modulating the evolutionary outcome overall in time-invariant environments and as a function of exposure epoch in time-dependent environments. This suggests that identifying strong selective agents and their pleiotropic effects is critical for predicting evolution in fitness seascapes, with ramifications for evolutionarily informed strategies to mitigate drug resistance evolution.

## Main

The genomes of evolving organisms can be portrayed as embedded in 3-dimensional fitness landscapes with low-fitness valleys and high-fitness peaks.^1,2^ The initial position on the landscape depends on the genomic background, which varies between clades within a population, and determines the mutations with positive fitness consequences. Once a mutation with a sufficient selection coefficient for the landscape position occurs, it will be selected, causing movement upwards from a valley or towards a peak. This position, in turn, determines the subsequent mutations with high selection coefficients. Ultimately, the population becomes trapped on a local peak or reaches a global peak depending on the starting position, traversed path and ruggedness of the landscape. The concept of a 3d fitness landscape is based on the realization that the fitness effects of mutations virtually always depend on the genomic background (pervasive epistasis). Typically, the topology of the landscape is presented as static. This may hold in a minimal setup with the evolution of a single gene in a stable environment and when complications such as frequency dependent selection are not present. What features of such static landscapes affect the predictability of evolution has become an active research field.^3,4^ However, in many realistic scenarios, the targets of selection change over time, causing the fitness landscape also to change over time, becoming a fitness seascape.^4,5^ For instance, mutations improving adaptive traits often have deleterious consequences on conserved traits, an example of pleiotropy, causing selection to alternate between mutations improving the adaptive trait and mutations compensating for the cost on the conserved trait. Moreover, the concept of fitness seascape inherently captures evolution along time-dependent selective environments.

In fitness seascapes, rather than the stepwise refinement of a single trait depending on a limited set of epistatic interactions, the mutational path is better characterized as a serpentine path where previous adaptations can have varied effects on where the genome lands in each time-dependent landscape.^4^ One consequence of this is that directional selection improving a particular trait is likely to be stronger for time-invariant compared to time-dependent environments. However, mutations improving an adaptive trait in a particular selective environment often influence other traits (*i*.*e*. display pleitropy), which may make the organism either more or less adaptive to a subsequent selective environment compared to the initial state. Pleiotropic effects have been shown to be prevalent in multiple systems, including microbes evolving antimicrobial resistance.^6^ Strong pleiotropic effects can have dramatic effects on the evolutionary outcome as key traits may improve even in the absence of direct selection.

In time-dependent environments, epistasis and pleiotropy are expected to give rise to historical contingency of evolution whereby the mutations that are adaptive in the current environment are contingent on the adaptive mutations accrued in the previous environment(s).^7-9^ This should be seen as differences in the mutational paths and profiles as well as in phenotypic outcomes when the order of the environments is altered, with some sequences constraining and others potentiating evolution. From a statistical viewpoint, the variance of outcomes should therefore be greater than expected by chance. A low deviation from the null expectation would indicate a negligible role for epistasis and pleiotropy in adaptation. For instance, in the case of strong selective pressures, only a single or narrow set of mutational targets may enable evolutionary rescue in each environment, causing the mutational profile and adaptive trait outcome in a time-dependent environment to be simply an aggregate of exposure to each environment in isolation. However, since strong adaptations often have deleterious consequences on conserved traits (among the most frequent modes of pleiotropy), accruing a succession of strong adaptations each accompanied by a fitness cost may prove deleterious after a particular set of environments. This could cause the extinction probability to increase as a function of the number of environments. A dependence of extinction dynamics on the environmental sequence, in turn, would indicate a stronger role for epistasis or pleiotropy.

Even though evolution in real-world systems typically occurs in dynamic fitness seascapes rather than static landscapes, evolutionary dynamics in fitness seascapes remain poorly understood. Among the underexplored questions feature the relative contribution of directional selection and pleiotropy on the evolutionary outcome in time-invariant versus -dependent environments. Moreover, the extent to which the environmental sequence determines the evolutionary outcome in time-dependent environments has not been comprehensively examined. The relative role of these processes influences the predictability of evolution and determines the conditions where adaptation is constrained or enhanced, which is key also for any practical efforts to guide evolution, such as in evolutionarily informed strategies to treat cancer or mitigate antimicrobial resistance.^7,10,11^

Here we designed a minimal setup for assessing evolutionary outcomes in fitness seascapes, comparing time-invariant and -dependent environments, and including among the latter all possible permutations of environmental orders. We utilized a microbial model system exposing *Escherichia coli* at high replication to four environmental epochs consisting of three different antimicrobials at subinhibitory levels and an antimicrobial-free environment (Figure 1). We also performed the full experiment in the presence of a bacteriophage to investigate the modulatory impact of an added layer of selective pressure, resulting in a total of 928 bacterial populations. Phages are also of interest by adding another, biotic stress to the system as well as representing an alternative type of antibacterial agent, with an increasing trend in real-life clinical applications of phage therapy. We phenotyped populations over time for adaptation to each selective pressure as well as phenotyping and whole-genome sequencing clones isolated from populations at the experimental end-point. To assess differences in evolutionary outcomes between fitness seascapes, we used a combination of analyses, including information theoretical and machine learning approaches. This enabled us to identify characteristics of fitness seascapes constraining or enhancing adaptation and influencing the predictability of evolution. Our findings have implications for evolutionarily informed strategies to manage populations, such as mitigating antimicrobial resistance evolution.

**Fig 1.**
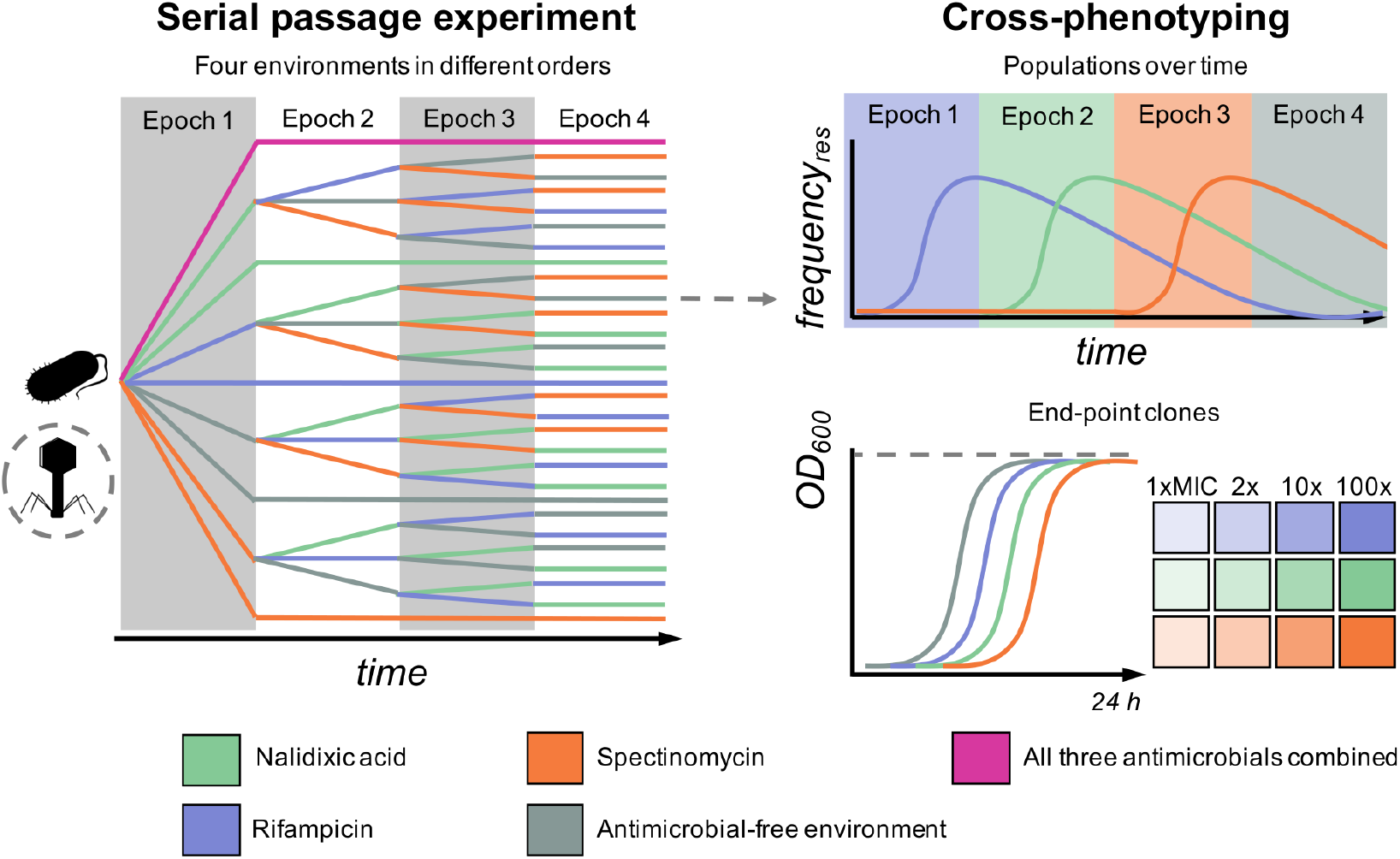
Overview of experimental design to systematically study evolution under fitness seascapes. The main experiment on the left was a 48-day serial passage experiment where initially isogenic *E. coli* was subjected to a control environment without antimicrobials; time-invariant environments with single antimicrobials or all three antimicrobials combined; and time-dependent environments encompassing all permutations of four 12-day epochs, including the three antimicrobials and one antimicrobial-free environment. The antimicrobials were used at subinhibitory concentrations (0.5 × MIC). The full experiment was repeated with and without initial introduction of phage representing an alternative type of selection pressure. Each unique treatment combination was replicated 16 times, amounting to a total of 928 independent populations, which were cross-phenotyped over time against low-level resistance to each of the antimicrobials. In addition, a single dominant clone was isolated from all surviving end-point populations (*N* = 900) and phenotyped for growth (optical density, OD, at 600 nm after 24 h culture) at several concentrations of each antimicrobial. To investigate the underlying molecular evolution, a subset of 235 clones representing different regimes and divergent low-level resistance outcomes were also subjected to whole-genome sequencing.

## Results

### Evolutionary dynamics and outcomes differ between and within time-invariant and -dependent environments

The evolutionary dynamics and outcomes differed both between and within time-invariant and -dependent environments (Fig. 2A–F; for statistical outputs for each selective agent at population and clone levels, see Tables S1—S6). Within the time-invariant environments, as expected, constant subinhibitory (0.5 × minimum inhibitory concentration, MIC) selection by a single selective pressure (*i*.*e*. antimicrobial agent) lead to the emergence of low-level (1 x MIC) resistance and phage selection to the emergence of phage resistance. Selection was strongest for one antimicrobial agent, rifampicin, with most replicate populations evolving low-level resistance both in the presence and absence of bacteriophage. The two other agents, nalidixic acid and spectinomycin, depended on the presence of bacteriophage for resistance to evolve to levels detectable for clones from the experimental end-point. Even then, low-level resistance only evolved in a minority of the replicate populations. The development of nalidixic acid and spectinomycin resistance in the presence of phage could be caused by a higher effective MIC (*i*.*e*. higher selection coefficient) in the presence of an additional stressor. More sensitive population-level phenotyping over time, where resistance in a subset of the population could yield a positive signal (while end-point clones represent dominant population phenotypes), showed stronger nalidixic acid resistance development, with a minor clade in most populations developing low-level resistance. This could not be assessed for spectinomycin as the temporal population-level data was too noisy (high levels of positive signal across environments). This is likely to be the outcome of random genetic variation in minor clades exhibiting low-level resistance phenotypes.

**Fig 2.**
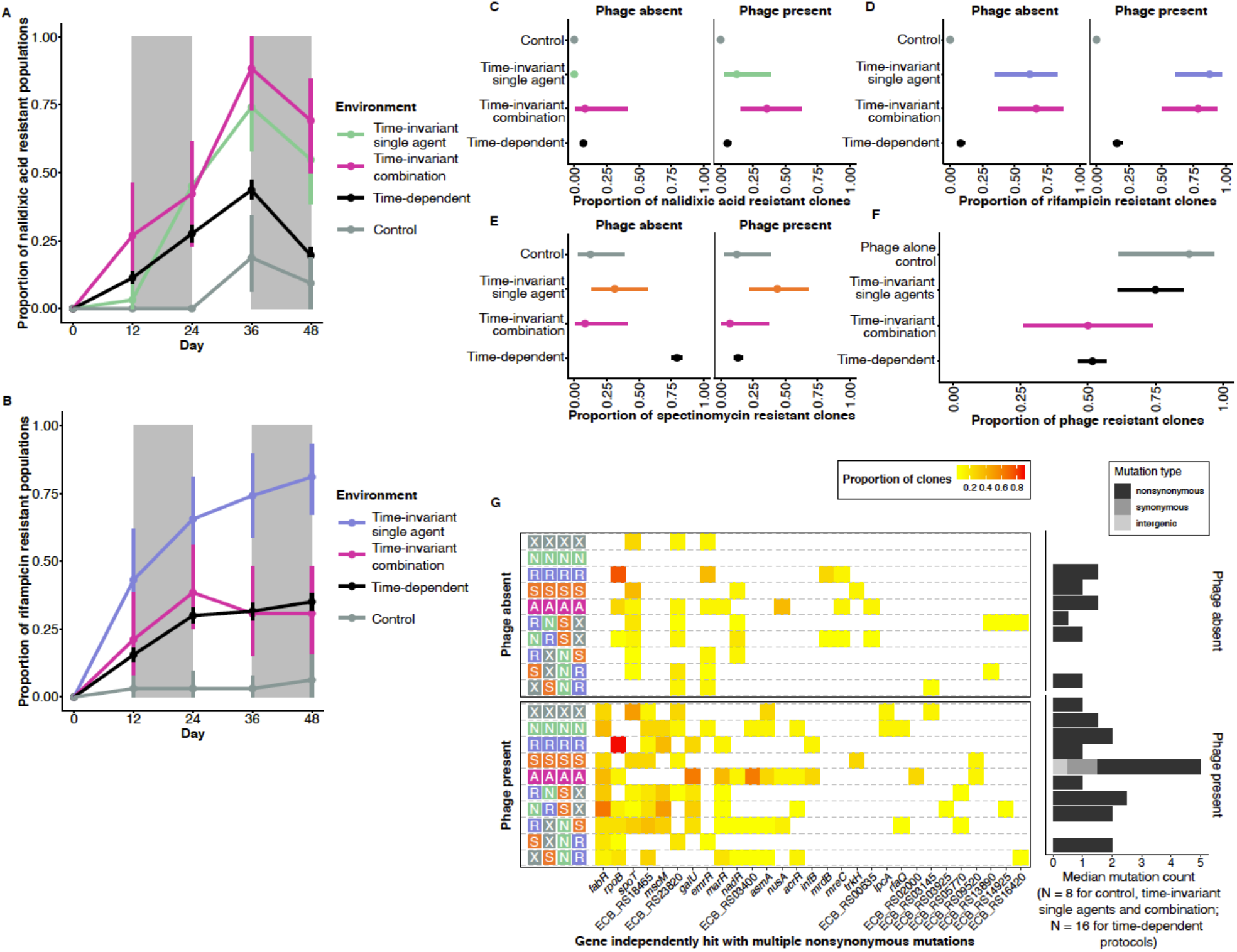
Contingency of phenotypic outcome and mutation landscape on selective regime. (A) and (B) show resistance dynamics over time for nalidixic acid and rifampicin, respectively (mean ± bootstrapped 95 % confidence intervals; 32 replicates per mean data point; data in presence and absence of phage pooled). Grey rectangles denote epoch boundaries for time-dependent protocols. (C), (D), (E) and (F) show low-level resistance outcomes to nalidixic acid, rifampicin, spectinomycin and phage T4, respectively, for clones isolated from each population at the experimental end-point (logistic regression expected value ± 95 % confidence intervals; 16 replicates per mean data point). Resistance has been quantified as a binary variable and indicates the ability to grow at levels exceeding the minimum inhibitory concentration of the ancestral bacterial strain. (G) Mutational landscape. The heat map on the left shows the proportion of sequenced clones containing a nonsynonymous (or infrequently synonymous) mutation in a gene recurrently hit in the dataset. The genes have been ordered by total number of hits. The bar plot on the right shows the median mutation count for clones in each history (lack of bar indicates median of 0 mutations). The *y*-axis labels indicate the antimicrobial therapy protocol with the following encoding for each of the four 12-day experimental epochs: X = antimicrobial-free environment; N = nalidixic acid; R = rifampicin; S = spectinomycin; A = all three antimicrobial compounds combined.

A time-invariant environment combining all three antimicrobials showed similar levels of rifampicin resistance and higher levels of nalidixic acid resistance (also in the absence of phage) compared to the single-antimicrobial environments, while spectinomycin resistance failed to evolve. The higher levels of nalidixic acid resistance could either be the outcome of the effective MIC being higher (*i*.*e*. higher selection coefficient) in the multidrug (*i*.*e*. multi-stressor) environment compared to the single-drug environment or pleiotropic effects from rifampicin resistance (inspected further below). Since spectinomycin exhibited the weakest selection pressure out of the three antimicrobials, the failure of spectinomycin resistance to evolve in the multidrug environment may be the outcome of the fitness cost of more readily evolved rifampicin or nalidixic acid resistance further decreasing the selection coefficient for spectinomycin resistance.

Evolution was strongly constrained for two out of the three antimicrobials, rifampicin and nalidixic acid, in the time-dependent compared to -invariable environments, with no to low proportions of replicate populations evolving low-level resistance. Spectinomycin was a striking exception, with no sign of resistance evolution in the presence of phage but high prevalence of resistance in the absence of phage (see next section for interpretation).

Phenotypic resistances to each of the antimicrobials as well as the phage were associated with recurrent mutations in particular genes (Fig. 2G). Some of these genes have previously been associated with phenotypic resistance to these agents, while others have not and are implicated with low-level phenotypic resistance in this study (for detailed information on these genes, see Supporting results). There were little signs of differences in recurrent genes between time-invariant and -dependent environments, although the low prevalence of resistance evolution in the latter prevents a thorough statistical testing of this question.

### Pleiotropic and fitness effects of strong selective agents, rifampicin and phage, modulate evolutionary outcome

Unexpectedly, the time-invariant rifampicin environment resulted in an increased probability of low-level resistance to nalidixic acid and spectinomycin, specifically in the absence of phage (Fig. 3A). When comparing control and time-invariant rifampicin environments with and without phage, nalidixic acid resistant clones only occurred in time-invariant rifampicin environments without phage (hence their prevalence between these environments cannot be compared using logistic regression). Low-level spectinomycin resistance, in turn, occurred in a small subset of the populations also in the control environments but was extremely prevalent in the time-invariant rifampicin environment without phage (time-invariant rifampicin environment vs.control, *P* = 0.039; phage presence, *P* < 0.001; interaction, *P* = 0.002; for full results, see Table S7). Moreover, low-level spectinomycin resistance was much more prevalent in time-dependent environments in the absence of phage compared to the time-invariant spectinomycin environment. We hypothesized that both observations could result from pleiotropic effects of mutations selected by rifampicin, with either mutational targets being altered or the pleiotropic effect being modulated by the presence of phage. In line with the latter explanation, we found that 6/8 clones containing *rpoB* mutations (producing rifampicin resistance) and unexposed to the phage displayed low-level resistance to spectinomycin. Conversely, only 4/17 clones with *rpoB* mutations in the presence of phage displayed resistance to spectinomycin. We also found a similar pattern of antimicrobial cross-resistance depending on phage resistance for two other genes: the stringent response gene *spoT* which was mutated across experimental treatments and the previously identified spectinomycin-selected gene *nadR* (Fig. 3B). Consequently, clones containing mutations in either *nadR, rpoB* or *spoT* had a high likelihood of exhibiting a low-level spectinomycin resistance phenotype conditioned on occurring in a phage susceptible genomic background (ANOVA for binomial generalized linear model: presence of mutations in *nadR, rpoB* or *spoT, χ*^2^_1,233_ = 1.90, *P* = 0.17; phage resistance, *χ*^2^_1,232_ = 25.7, *P* < 0.001; presence of mutations in *nadR, rpoB* or *spoT* × phage resistance, *χ*^2^_1,231_ = 7.01, *P* = 0.008).

**Fig 3.**
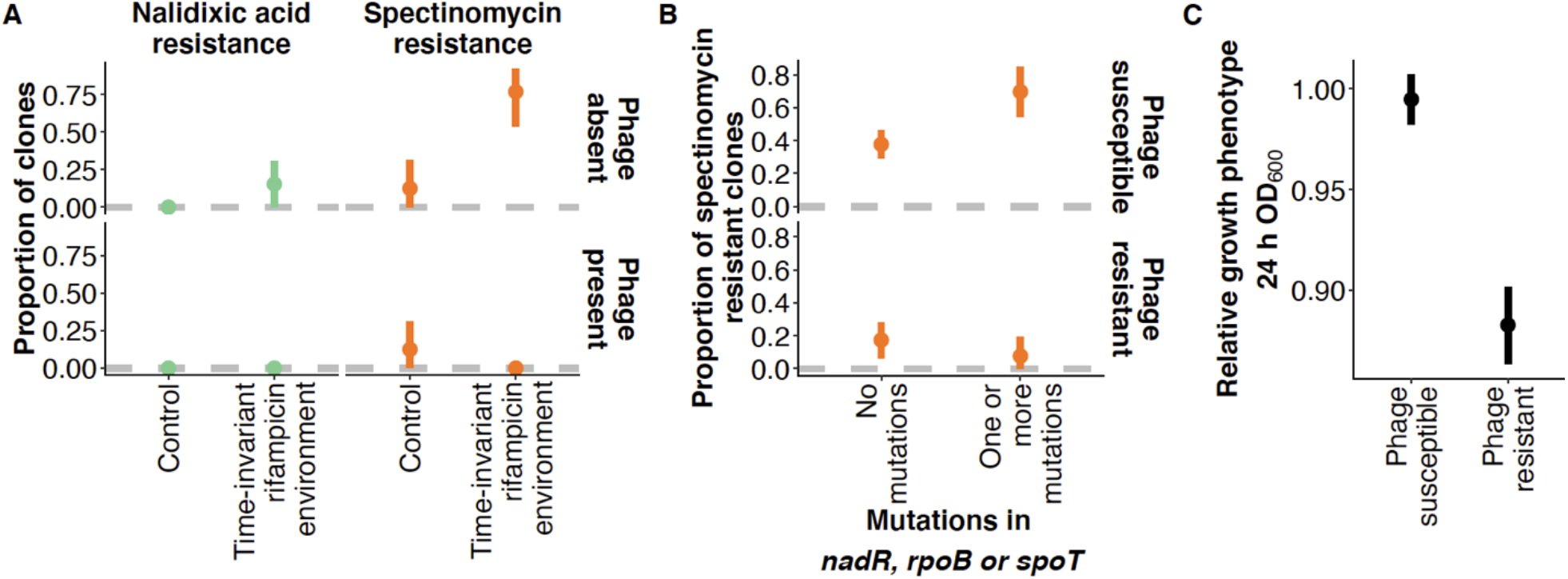
Pleiotropy and fitness effect of resistance. (A) Low-level nalidixic acid and spectinomycin resistance after 48 days in time-invariant rifampicin environment (mean ± bootstrapped 95 % confidence intervals). (B) Influence of phage resistance on whether mutations in the genes *nadR, rpoB* or *spoT* produce a low-level spectinomycin resistant phenotype (mean ± bootstrapped 95 % confidence intervals). The data is for clones from phage-exposed environments (for which phage resistance phenotype was determined). (C) Fitness effect of phage resistance (mean ± bootstrapped 95 % confidence intervals). Fitness has been quantified as optical density (OD) at 600 nm wavelength after 24 h culture in liquid medium. The value has here been related to the mean growth of the clones from the control treatment (absence of antimicrobials and phage). The data for all the figures is for clones isolated from populations at the experimental end point (*N*_*total*_ = 900), with subset treatments or phenotypes included in a particular analysis indicated in the figure or legend.

Furthermore, we found that all *mreC* mutants (selected by rifampicin) as well as a single *mreD* mutant displayed low-level multidrug resistance. Two of three *mreC* mutants (containing the same frameshift mutation, Glu291fs) and the *mreD* mutant were resistant to all three drugs, while a single *mreC* mutant (Val46Gly) was resistant to both nalidixic acid and spectinomycin but remained susceptible to rifampicin. Mutations in *mreC* and *mreD*, whose products work in concert to determine cell shape and elongation, did not occur in the presence of phage. In turn, mutations in *marR* selected by rifampicin that occurred only in the presence of phage also resulted in nalidixic acid resistance. These mutations could therefore explain the increased probability of nalidixic acid resistance in the presence of rifampicin. In addition, we found that phage resistance, associated, in particular, with mutations in *fabR, galU* and ECB RS18465, had a strong fitness cost as indicated by reduced bacterial growth after 24 culture (88.8 % growth of phage sensitive clones) in the absence of the phage or antimicrobial compounds (ANOVA for linear model: phage susceptibility, *F*_1,898_ = 84.1, *P* < 0.001) (Fig. 3C). Together these observations suggest that nalidixic acid and spectinomycin resistance dynamics were to a large extent driven by the other two selective agents (rifampicin and phage) imposing stronger selection through the following three mechanisms: i. cross-resistance (*i*.*e*. pleiotropic) mutations selected in the presence of rifampicin; ii. the effect of phage on the strength of selection for antimicrobial resistance and the targets of antimicrobial resistance mutations; iii. the loss of the antimicrobial resistance phenotype of specific mutations in a phage resistant background. Cases ii. and iii. may be related to the strong fitness-impairing consequence of phage resistance.

### Modifying effects of strong selective environments largely explain differences between time-dependent environments

The evolutionary dynamics described above largely determined differences in low-level resistance levels between the time-dependent environments. First, rifampicin which was the strongest selective antimicrobial compound caused resistance to occur as a function of exposure epoch mainly by selecting for *rpoB* mutations followed by negative selection post-exposure (Figs 4A & S1B; rifampicin exposure epoch, *P* = 0.074; for full results, see Table S8). Second, nalidixic acid resistance occurred at a much lower level in general (Figs 4B,C & S1A). It occurred as a function of rifampicin exposure in the absence of phage where selection by nalidixic acid was weak and rifampicin selected for low levels of nalidixic acid cross-resistance by *mreC* mutations (Figs 4B & S1A; nalidixic acid exposure epoch, *P* = 0.48; rifampicin exposure epoch, *P* = 0.003; for full results, see Table S9). In the presence of phage, however, nalidixic acid resistance was more strongly driven by nalidixic acid exposure selecting for *acrR* and ECB RS03400 mutations (Figs 2G, 4C & S1B; nalidixic acid exposure epoch, *P* = 0.007; rifampicin exposure epoch, *P* = 0.22; for full results, see Table S10). This is consistent with the bacterial cells experiencing stronger selection for nalidixic acid resistance in the presence of both phage and nalidixic acid compared to nalidixic acid alone. Finally, in line with cross-selection by rifampicin, spectinomycin resistance level was influenced by both the spectinomycin and rifampicin exposure epochs, although being mainly determined by the presence of phage (Figs 3B & S1C; spectinomycin exposure epoch, *P* = 0.012; rifampicin exposure epoch, *P* < 0.001; phage presence, *P* < 0.001; for full results, see Table S11).

**Fig 4.**
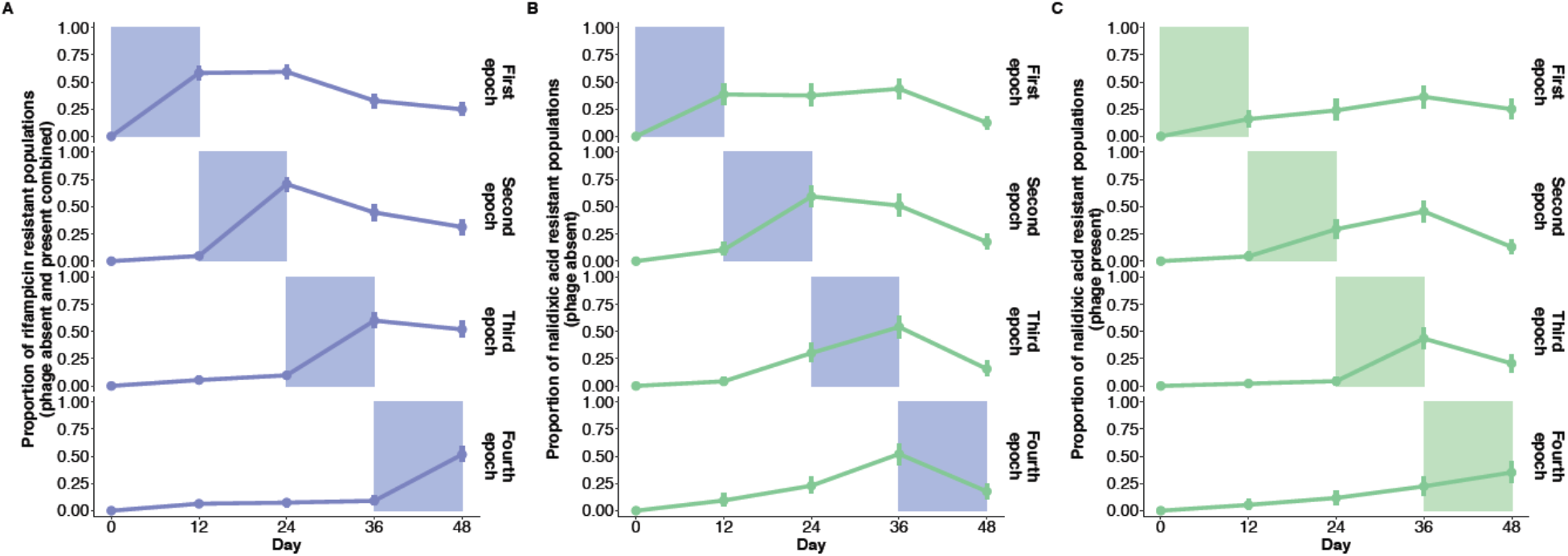
Antimicrobials driving resistance evolution in time-dependent regimes. (A) Rifampicin resistance over time as a function of rifampicin exposure epoch (both in presence and absence of phage which had no effect on selective antimicrobial as it did for nalidixic acid). (B) Nalidixic acid resistance over time in the absence of phage as a function of rifampicin exposure epoch. (C) Nalidixic acid resistance over time in the presence of phage as a function of nalidixic acid exposure epoch. All the data is shown as mean resistance ± bootstrapped 95 % confidence intervals, and is based on *N* = 928 populations. The shaded area indicates the relevant (antimicrobial color code) exposure epoch.

A high prevalence of low-level spectinomycin resistance in the absence of phage accounted for a large proportion of positive resistance signals in the phenotypic data set. Because of this, low-level multidrug resistance (here referring to resistance to multiple antimicrobial compounds and excluding resistance to the phage) was more likely to occur in the absence of phage despite the phage exacerbating selection for rifampicin and nalidixic acid resistance in the time-invariant single-drug environments. As nalidixic acid selection was weak, most cases of multidrug resistance were cases of rifampicin-spectinomycin cross-resistance. Notably, three out of five among the sequenced clones displaying resistance to all three agents contained mutations in the cell-shape determining genes *mreC* and *mreD*. Although rifampicin resistance levels began to decay after the rifampicin exposure epoch, likely owing to a fitness cost of rifampicin resistance, the effect was too weak to introduce a clear history dependence effect on low-level multidrug resistance levels. Therefore, differences in the evolutionary outcome between the time-dependent environments were mostly accounted for by the following factors: differences in selection strength between the agents; pleiotropic effects of rifampicin resistance; and modifying effects of phage exposure on antimicrobial resistance evolution.

### Inference of environmental past and predictability of evolutionary future are strongest for driver environments

The strong selective environments (rifampicin and phage) largely accounting for the low-level multidrug resistance landscape, correspondingly, exhibit the strongest predictive power regarding the past drug exposure and future resistance outcome of the bacterial populations (Fig. 5A). As expected based on the strong fitness consequence of phage resistance, machine learning (random forest) models were able to predict past phage exposure with high accuracy based on growth data from end-point clones in the absence or presence of different levels of the experimental antimicrobials. The same data could also be used to train a model to precisely predict the rifampicin exposure epoch, consistent with rifampicin resistance levels both decaying after exposure and influencing the overall resistance phenotypes. Conversely, in line with expectations, high-accuracy predictive models could not be constructed for the exposure epoch of the weak selective agent nalidixic acid or cross-selected agent spectinomycin. These factors were also seen as modest predictive power of models predicting the full antimicrobial exposure sequence (*i*.*e*. environmental past). Therefore, the ability to predict the environmental past from the current phenotypic state is increased by high selection coefficients and strong fitness effects of resistances and can be obscured by low selection coefficients and pleiotropy.

**Fig 5.**
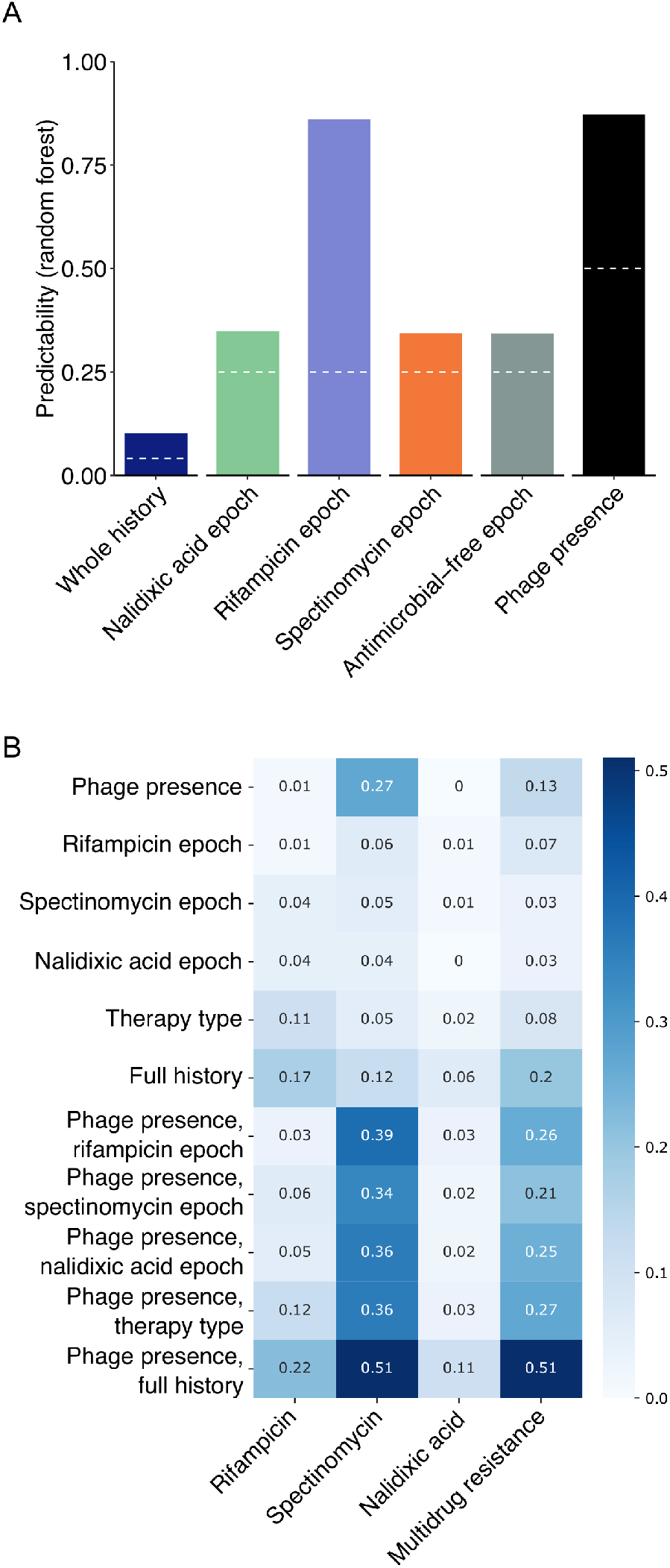
Predictability of past drug exposure and future resistance evolution in time-dependent regime. (A) Degree of predictive power obtained for antimicrobial and phage exposure past based on phenotypic data from populations and clones at the end-point of 48-day serial passage experiment. The white dashed line indicates the level of predictability of the estimated factor by chance. (B) Mutual information between exposure history (rows) and end-point clone antimicrobial resistance phenotype states (columns). The exposure histories are described at various levels of detail ranging from coarse (phage status and epoch of exposure to an antimicrobial separately) to full information (phage status and antimicrobial epoch order considered simultaneously). For example, the phage presence label of the histories carries substantial information on spectinomycin resistance status of the end points clones.

We further quantified information theoretically the relationship between exposure histories and each of the end point phenotypes individually by evaluating their mutual information (Fig. 5B). A large value of mutual information between an environment and a phenotype indicates that knowing one substantially removes uncertainty about the other. Phage and spectinomycin exhibited strong mutual information (0.27 std 0.03) and adding the detail of the antimicrobial exposure order further increased it to 0.51 std 0.03. For nalidixic acid and rifampicin, knowing whether the exposure history had phage or not carried little information. However, for both compounds, increasing the detail of the antimicrobial exposure order carried information. In contrast to the random forest modelling results, we noticed that the epoch of exposure to rifampicin did not greatly reduce uncertainty for the individual end-point resistance states. This is due to the random forest model exploiting both population and clone data at multiple MIC values – beyond binary (quantitative OD value instead of 0 for susceptible and 1 for resistant) – as well as using the joint phenotype, respect to all compounds, as its basis for predicting the past exposure environment.

Together these results are consistent with the previously reported strongest environment-phenotype links, namely, the relationship between rifampicin and phage exposures and corresponding resistances, and the (inverse) relationship between phage presence and low-level spectinomycin resistance. Therefore, establishing the relationship between driver environments (here, rifampicin and phage) and phenotypic states is critical for understanding and predicting evolution along time-dependent fitness seascapes.

## Discussion

Inspecting differences in evolutionary dynamics between time-invariant and -dependent fitness seascapes, we found that specific environments drove environment-phenotype links. These driver environments (here, one subinhibitory antimicrobial and a phage) were characterised by strong selective pressure on the target adaptive trait and pleiotropic effects on off-target traits. Furthermore, we showed that establishing the driver environments and their phenotypic consequences is key for predicting evolution along fitness seascapes. Our data is also consistent with the order of the driver agents strongly influencing the evolutionary outcome in time-dependent fitness seascapes compared to the order in general. The practical importance of this finding is stressed by the fact that time-dependent fitness seascapes in real-life scenarios are highly unlikely to contain a set of environments equivocal in terms of the strength of selection and the strength of epistatic and pleiotropic effects. Therefore, any effort to construct general laws and models for evolution along fitness seascapes should account for temporal or environmental differences in the strengths of these factors.

Although our experimental setting utilizing sub-MIC selection (0.5 × MIC) did not mimic therapeutic antimicrobial levels, our findings have a number of potential implications concerning the global antimicrobial resistance crisis. Firstly, sub-MIC levels of antimicrobials occur in many human-impacted environments such as wastewater and agricultural environments, as well as in the concentration gradients within the tissues of medicated humans, production animals and pets. In these conditions, depending on the antimicrobial compound, even very low concentrations can increase the fitness of resistant cells above that of susceptible cells, causing positive selection for resistance.^12^ Because of weaker selection compared to high antibiotic levels, mutations producing resistance in such conditions cannot afford to be coupled with strong fitness costs. This can lead low antibiotic level environments to facilitate the stepwise evolution of high-resistance, low-fitness-cost, mutants particularly problematic to remove if they spread among humans or production animals as pathogens (including opportunistic pathogens).^13^ Our findings demonstrate that particular driver agents can create low levels of resistance against multiple drugs even at sub-MICs, a phenomenon (pervasive pleiotropy causing cross-resistance or collateral susceptibility) studied previously mainly for high antimicrobial levels.^6,14,15^ This finding expands the scope of the potential undesired consequences of environmental antimicrobial residues. Such environments are also highly likely to experience residues of different antimicrobials over time, creating fitness seascapes similar to those included in our study setup. Under such conditions, it may be important to establish the historical exposure of the environment to particular driver agents as part of risk estimation for subsequent antimicrobial contamination.

Second, the driver agents modified the interplay between selection and the epoch length by exacerbating selection and thus partially removing the desired filtering effect of epochs to resistance evolution. As the driver agents affected resistance evolution to both directions, assessing their overall impact for a specific therapy requires experimentation. Clearly, identifying and testing the impact of such driver agents has potential for therapy optimization and their efficient usage should be studied further using eco-evolutionary control theory.^16^ Intriguingly, driver agents do not necessarily need to be antimicrobials, for instance, the effects of stress environments can be further modulated by inhibiting global regulators.^17^

Interestingly, it was recently demonstrated that cellular hysteresis whereby transgenerational changes in cellular physiology induced by one drug alter the bacterial response to a second drug can influence bacterial evolutionary trajectories in alternating drug protocols, particularly with rapid (every 12 or 24 h) switching.^18^ It is conceivable that cellular hysteresis could have contributed to the evolutionary outcomes in this study. Since the number of putative resistance mutations accumulated over exposure to three drugs and phage tended to be low (two on average), pleiotropic effects of previously accumulated mutations are likely to explain only a fraction of the variance in evolutionary trajectories between time-dependent protocols. An interesting avenue of future exploration is whether the driver environments are also more likely to induce cellular hysteresis. This could contribute to particular drugs functioning as driver agents in multidrug systems.

Our study also lends general insights into ecological resilience critical to understand owing to the climate and biodiversity crisis. Based on our study findings, when organisms evolve in temporally changing environments, particular environments (especially stressors) are likely to play a pivotal role in facilitating or obstructing multi-environmental adaptation relative to the sequence of environments alone. Identifying such key environmental conditions and their off-target effects may be critical in fields such as nature conservation for preventing population or community collapse and for enhancing their biological resilience. Our study and previous studies with microbial systems suggest that strong off-target effects of adaptations to particular environments may be commonplace. Therefore, a failure to incorporate them in ecological and evolutionary predictions can severely restrict the efficacy of interventions based on them.

Finally, fitness seascapes have been studied both experimentally and theoretically much less than their static counterparts. A reason behind this gap are the apparent complexities involved - it is much harder to convince oneself that a particular fitness seascape constitutes a minimal model worth studying at depth compared to the canonical models of static landscapes familiar from textbooks^19^ However, our results show that experimental work on the topic is both feasible and informative for future theoretical work. Indeed, the possibility of simplifying evolution under complex seascapes by describing them in terms of few a driver agents looks theoretically tractable.

## Methods

### Model organisms

As a model organism, we used the *E. coli* strain REL606 (Ara^−^) obtained from the Yale Stock Center (*E. coli* B ATCC 11303). To enable distinguishing strains from mixed cultures in potential competition assays, we produced a REL607-like (Ara^+^) mutant by culturing REL606 on minimal arabinose plates (personal correspondence with Richard Lenski). The mutant was verified by Sanger sequencing by a third party (Institute of Biotechnology, University of Helsinki), and has the same point mutation as REL607, allowing the use of arabinose as a carbon source and thereby chromogenic differentiation from REL606 on arabinose agar (Fig. S2). A co-culture competition assay was used to determine that the two strains did not initially differ in fitness (Fig. S3). Additional details for mutant isolation and the competition assay are described in the Method Details and Tables S12, S13 and S14. As virulent bacteriophage, we used T4 strain ATCC 11303-B4. All culturing steps across experiments were performed at 37 °C.

### Serial passage experiment

We conducted a 48-day serial passage experiment consisting of four 12-day epochs (Fig. 1). To allow an exhaustive exploration of equivalent antimicrobial exposure space, we subjected initially isogenic populations of *E. coli* to all permutations of four antimicrobial treatments (one epoch each). This generated a total of 24 unique exposure histories containing the same antimicrobial treatments in different orders. In addition, we included five sequences, three of them featuring the same antimicrobial treatment (*i*.*e*. time-invariant single agent environment) and one containing all antimicrobials combined (*i*.*e*. time-invariant combination environment) across all epochs throughout the experiment, as well as an antimicrobial-free control environment. The four antimicrobial treatments for the four 12-day epochs in the permutation (*i*.*e*. time-dependent) environments consisted of three antimicrobial-containing and one antimicrobial-free treatment. The antimicrobial-free treatment was added since antimicrobial-free periods can have a major effect on resistance dynamics by either reversing prior resistance owing to fitness costs or by potentiating future adaptation through compensatory adaptations or increased genetic heterogeneity. The three antimicrobials were selected based on the previously established susceptibility of *E. coli* REL606 and its ability to *de novo* evolve resistance to them, as well as difference in antimicrobial class, mode of action and genomic target of resistance mutations. Different antibiotic classes were used to avoid major synergy or antagonism. However, since such effects at weaker levels are extremely common, we considered that they cannot be completely ruled out and did not screen for them at the experimental design phase. The antimicrobials thus selected were nalidixic acid (naphthyridone, quinolone-like antimicrobial targeting DNA gyrase), rifampicin (rifamycin antimicrobial targeting RNA polymerase) and spectinomycin (aminocyclitol, aminoglycoside-like antimicrobial targeting 30S subunit of ribosome). In addition to the antimicrobials, the full experiment was performed both with and without the bacteriophage T4, representing an alternative selective pressure or drug type (phage therapy) with a likely even more divergent cellular target and mode of resistance compared to those between the antimicrobial compounds.

The experiment was performed in a deep 96 well plate setup in the DM1000 medium, which produces approximately 2 × 10^9^ cells mL^−1^ during a 24 h culture cycle at 37 °C. The experiment was started by adding approximately 10^6^ cells from a 24 h culture of *E. coli* to a final volume of 500 µL of DM1000 containing the appropriate antimicrobials. The antimicrobials were used at 0.5 × minimum inhibitory concentration (MIC) to cause relatively strong selection for resistance while not immediately killing susceptible cells or causing extinction of the bacterial population in the presence of phage due to synergistic population crash. For the phage treatments, 5 × 10^5^ infective particles (constituting a multiplicity-of-infection value, or MOI, of 0.5) were subsequently added to the wells. Antibiotic MIC and phage MOI are not comparative measures as virulent phages are multiplying entities that kill susceptible bacteria while antibiotic MIC is a static concentration which kills susceptible bacteria only at concentrations starting from the MIC level. Both phage MOI and antibiotic level relative to MIC affect resistance evolutionary dynamics in important ways. The lower the MOI, the more generations bacteria have to evolve resistance until the majority of the population will encounter the phage. Similar to low MOI levels, sub-MIC levels of antibiotics that are high enough to cause positive selection for resistance do not require resistance to be immediately present in the standing genetic variation of the population but resistance may evolve over time in a susceptible population. The well plates were cultured at 37 °C with constant shaking at 120 r.p.m.

Every 24 h, 2 % (10 µL) of the old culture was transferred to a new well containing fresh medium and the appropriate antimicrobial. However, the phage was allowed to go extinct without replenishment. Every 96 h, the cultures were freeze-stored with glycerol at –20 °C for later analyses. Details concerning antimicrobial MIC determination using the microdilution method, the culture medium and antimicrobial concentrations can be found in Method Details sections below. To ensure adequate statistical power, each of the unique treatment combinations (N = 58, including 24 antimicrobial sequences and five monotherapy, combination therapy or control sequences in absence/presence of T4 phage) was repeated 16 times, eight times each for the REL606 and REL607-like strain. This resulted in a total of 928 unique serially passaged *E. coli* populations.

### Measuring antimicrobial resistance phenotypes

To quantify the evolution of antimicrobial resistance over time, every 2 days in the 48-day evolutionary experiment, 2 % (10 µL) of the old culture was transferred, using a 96-pin replicator, to a large petri dish containing lysogeny broth (LB) agar supplemented with one of the three antimicrobials at a selective concentration (> 1 × MIC), including an antimicrobial-free plate to control for bacterial extinction. After culturing for 24 h at 37 °C, growth on the plates was quantified on a binary scale (0 = no growth; 1 = growth). Therefore, for this metric, growth indicates the detectability of antimicrobial resistance in a minimum of 2 % of the bacterial population. Notably, we cross-phenotyped all populations from all treatments and time points against all antimicrobials. Since the temporal tracking of resistance evolution employed a coarse measure diverging from experimental conditions (solid medium vs. liquid medium; higher antimicrobial level compared to selective concentration of 0.5 × MIC; phenotype based on fraction of heterogeneous population), we isolated individual clonal lineages from the experimental end-point and phenotyped them more precisely. Clones were isolated by streak-plating inoculum from the population on an LB agar plate, culturing overnight, and picking an individual colony, followed by propagation in liquid medium. The clones were phenotyped by culturing a small initial number of cells for 24 h in DM1000 liquid medium without antimicrobials or supplemented with one of the three antimicrobials at a range of multiplicities of the MIC value of the ancestral *E. coli* strain: 1, 2, 10, or 100 × MIC. This allowed obtaining quantitative growth and antimicrobial resistance phenotypes (in units of optical density at 600 nm, OD_600_) in the experimental conditions for isogenic lineages likely to represent the dominant *E. coli* genotype in each of the 900/928 populations that escaped extinction during the 48-day serial passage experiment. Before downstream analyses, the OD data was curated by removing background OD values specific to 96-well plate location, drug type, and drug concentration (Fig. S4), and following this, by removing (transforming to zero growth) likely false positive values based on belonging to a distinct distribution close to zero (Fig. S5). The cut-off used to remove false positives was 3 times the standard deviation of the noise distribution close to zero, representing a demarcation region between the noise distribution and true positive value distribution. In later analyses, resistance was defined for the clones as the capacity to grow at antimicrobial concentrations exceeding the MIC of the ancestral strain, as the experimental concentration of 0.5 × MIC selected mostly for low-level resistance (Fig. S6).

### Measuring phage presence and resistance phenotypes

We also measured the loss of phage from phage-containing populations during the experiment using the population level pin replicator system on agar plates containing a soft agar lawn of the ancestral *E. coli* strain, where the formation of a plaque indicates presence of phage. Moreover, we quantified the phage resistance phenotype of the end-point clones using a similar setup, with a lawn of the ancestral T4 phage instead of the bacterium. Bacterial growth on the phage lawn was taken to indicate phage resistance, which was therefore characterized as a binary trait (susceptible/resistant). Detailed phenotyping protocols are documented below.

### Plaque assay and generation of T4 phage stock

The T4 phage stock for the experiment was generated based on previously established protocols (Lenski lab protocol: lenski.mmg.msu.edu/ecoli/phagepharm.html, accessed 2018-05-15; Barrick lab protocol: barricklab.org/twiki/bin/view/Lab/ProceduresPhageFarming, accessed 2018-05-18). First, a dilution series of an existing T4 stock was prepared in LB liquid medium, and 100 µL was combined with 100 µL of exponentially growing host strain (REL606). Following incubation at 37 °C for 30 min, the mixture was added to 4 mL of soft LB agar (7 g agar per litre) kept at 55–60 °C, mixed and poured on top of an LB agar plate. The plates were cultured for 24 h at 37 °C, generating phage plaques at dilutions where the plaques are not touching. Subsequently, 1 mL of overnight host culture was transferred to 4 mL of LB medium in a 50 mL flask, and a toothpick was stuck into an isolated plaque and dropped into the host culture, which was kept at 37 °C with constant shaking at 150 r.p.m. After 6–8 h causing complete lysis of the host, 10 drops of chloroform were added and mixed to kill the remaining bacteria, and cell debris was removed by spinning down and keeping the supernatant. The phage was then titred as described above to determine phage density in the stock (as plaque forming units, or PFU, *i*.*e*. infectious phage particles). The lysate was stored with glycerol at –80 °C.

### DNA extraction, sequencing, and pre-processing sequence data

DNA extraction for clones from the experimental end-point was performed using the Qiaqen DNeasy 96 Blood & Tissue kit according to a custom protocol (detailed protocol below). DNA concentration was measured with the Qubit^®^ 3.0 fluorometer (Thermo Fisher Scientific, Waltham, MA, United States) using the HS assay kit. Whole genome sequencing was performed with the Illumina NovaSeq SP 300 (2 × 150 bp) platform by a third party (Institute for Molecular Medicine Finland, FIMM) according to in-house protocols. FASTQ files obtained from FIMM were quality controlled with Cutadapt v1.10 ^20^, including removal of sequencing adapters (with minimum overlap, -O, of 10 bp set for adapter match) and trimming sequences by allowing minimum Phred-scaled quality cutoff (-q) of 28 for the 30 end of each read, and minimum length of 30 bp. Before and following quality control, the quality of the sequence data was assessed with FastQC v0.11.8 (http://www.bioinformatics.babraham.ac.uk/projects/fastqc) and MultiQC v1.7 ^21^.

### Genome alignment, variant calling, and annotation

Quality controlled FASTQ files were aligned to the reference genome (NCBI Reference Sequence NC 012967, assembly ASM1798v1) with Bowtie 2 v2.3.4^22^ using default settings. SAMtools v1.4^23^ was used to convert thus obtained SAM files to BAM files, and to sort and index BAM files. Picard v2.18.10 (http://broadinstitute.github.io/picard) was used to mark duplicates and add read group information to BAM files, and following these steps, to compute genome coverage and other alignment metrics (commands CollectAlignmentSummaryMetrics and CollectWgsMetrics). Variant calling was performed with the Genome Analysis Toolkit (GATK) v3.8^24^, including indel realignment (commands RealignerTargetCreator and IndelRealigner), and variant calling for each clone separately with a combination of HaplotypeCaller (sample ploidy set to 1) and GenotypeGVCFs. Subsequently, the GATK commands SelectVariants and VariantFiltration were used to hard filter single-nucleotide polymorphism (SNP) and short insertion and deletion (indel) data separately, using the following criteria: for both, the absence of any variants detected in the ancestral strains (–discordance), a minimum Phred-scaled quality *P*-value normalized by allele depth (QD) of 2 to remove low-quality variants, and a maximum of 2 variants in a 20 bp window (–clusterSize 2 and –clusterWindowSize 20) to remove potential variant dense regions; for SNP data, a maximum Phred scaled *P*-value from the Fisher’s Exact Test to estimate strand bias (FS) of 60, a minimum root mean square mapping quality over all the reads at the site (MQ) of 40, a minimum value of –12.5 from the Rank Sum Test mapping qualities comparing the reads supporting the reference allele and the alternate allele (MQRankSum), and a minimum value of –8.0 from the Rank Sum Test for site position within reads (ReadPosRankSum); for indel data, maximum FS of 200, and minimum ReadPosRankSum of –20.0. Filtered genomic variant data was annotated against the reference genome with SnpEff v4.3i^25^. The ancestral REL606 strain used in the experiment had no variants relative to the reference genome, while the REL607-like strain was confirmed to have the *araA* mutation as well as one additional variant, a stop-lost variant in the pseudogene ECB RS25640 encoding a hypothetical protein.

### Regression and machine learning analyses

The regression and machine learning analyses (Figs 2A–F, 3, 4 and 5A) were performed in the R v3.6.1 environment^26^. Binomial generalized linear (*i*.*e*. logistic regression) models, with binary drug resistance or phage resistance outcome as a response variable and antimicrobial regime (time-invariant single agent, time-invariant combination or time-dependent protocol), antimicrobial exposure epoch, phage presence and presence of nonsynonymous mutations in genes of interest as covariates, were performed for experimental end-point populations or clones using the glm function in base R. Time series resistance data for time-invariant nalidixic acid and rifampicin environments were analyzed using generalized least squares models, as implemented in the *nlme* package^27^, assuming AR1 residual correlation structure within replicates. Random forest models for predicting exposure epochs from end-point phenotypes were generated using the *randomForest* package^28^. Before analyses, control treatments were excluded and rare features were removed based on min. 90 % of values being zero, and the data was standardized by converting each value into a *Z*-score (subtracting each sample’s mean and dividing by the sample’s standard deviation). The package *splitstackshape*^29^ was used to assign 80 % of each antimicrobial history into a training set and the rest 20 % into a test set. This was followed by random forest classification using the function randomForest implementing the Breiman’s random forest algorithm, with the options importance = TRUE and proximities = TRUE. The model was evaluated on the test set using the predict function, and the observed and predicted data were used to produce a confusion matrix with the confusionMatrix function in the *caret* package^30^. To avoid prediction bias due to a small test set size, this procedure was iterated 100 times, and the resulting confusion matrices were averaged over the iterations to produce one aggregate confusion matrix. The predictability of each estimated factor was interpreted as the weight of the diagonal (observed value matches predicted value) in the confusion matrix.

### Mutual information analyses

The analyses were performed in Python, using NumPy and pandas. We first bootstrapped 10,000 individual datasets drawn from the full dataset. Subsequently, the mutual information between experiment outcome and experimental conditions (*i*.*e*. environmental history) was calculated on these sets and averaged to a single value.

The following definition of mutual information was used for the calculation:

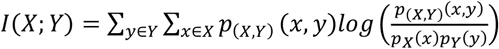

The marginal probabilities *p*_*X*_(*x*) were calculated by counting the number of samples corresponding to every outcome *x* – the level of resistance to the antimicrobial of interest, or the sum of these resistances as in the *multidrug resistance* column in Fig. 5B – and dividing the counts by the total number of samples.

The marginal probabilities *p*_*Y*_(*y*) were calculated by counting the number of samples which were subjected to experimental conditions *y*, and dividing the counts by the total number of samples; these conditions correspond either to: exposure to phage; the first epoch a particular antimicrobial has been administered if at all; the type of treatment used, grouping the alternating treatments together while considering the uniform treatments individually; and finally, the full exposure histories. These conditions were then further distinguished additionally by whether the phage had been administered or not. Similarly, the joint probabilities *p*_*X,Y*_(*x,y*) were calculated by counting the number of samples corresponding to every outcome *x* while subjected to experimental conditions *y*, and dividing the counts by the total number of samples.

## Data and code availability

Raw sequence read data has been deposited in the NCBI SRA under the accession PRJNA768654. All code and pre-processed data needed to reproduce the downstream analyses and figures will be available via Dryad/GitHub.

## Acknowledgments

We thank Roosa Jokela and Jutta Kasurinen for technical assistance; Chris Illingworth for comments on an earlier version of the manuscript; the group of Leopold Parts in the Wellcome Sanger Institute, UK, for hosting JC as visiting worker in 2019–2020; and CSC – IT Center for Science for the allocation of computational resources.

## Author contributions

Conceptualization: JC, TH, VM. Supervision: JC, VM. Data curation: JC, TM, FB, VM. Formal analysis, visualization and investigation: JC, VM, FB, TM. Writing – original draft: JC. Writing – review & editing: all authors.

