## Supplementary Information for "Strong selective environments determine evolutionary outcome in time-dependent fitness seascapes"

**This PDF file includes:**

Supplementary methods

Confirmation of REL607-like Ara+ reversion mutation

Competition assay for ancestral strains: *E. coli* REL606 and REL607-like Ara+ reversion mutant

Details on serial passage experiment

Determining minimum inhibitory concentrations for antimicrobials for ancestral *E. coli* strain

Details on phenotyping experimental populations over time

Details on phenotyping clones isolated from experimental end-point

Details on extraction of DNA from clones isolated from experimental end-point

Supplementary results

Details on potential molecular targets of resistance

Figures S1 to S7

Tables S1 to S14

### Supplementary methods

**Confirmation of REL607-like Ara+ reversion mutation**

The revertant produced for this study contains the same ara mutation (92G / GGC; REL606 ancestor: 92D / GAC) as the REL607 strain used in the *E. coli* Long-term Experimental Evolution project in the Lenski lab (barricklab.org/twiki/bin/view/Lab/ProtocolsAraMarker, accessed 2018-05-15). The *E. coli* B strain REL606 was acquired from the *E. coli* Genetic Resources at Yale CGSC, The Coli Genetic Stock Center (REL606 CGSC#: 12149). A liquid culture of the strain was streak-plated on Lysogeny Broth (LB) agar medium, and five clones were isolated by culturing five separate colonies in DM1000 medium and freeze-storage with glycerol at –80 ºC. All culturing steps were performed for 24 h at 37 ºC (liquid cultures with constant rotation at 150 r.p.m.), unless specified otherwise. To select for REL607-like Ara+ revertants derived from REL606 (Ara–), a colony of each of the five clones was cultured in six replicates of 10 mL of DM1000 medium, spun down to concentrate cells ($10^9$ cells), and plated on MA agar medium, followed by culturing for 48 h (personal correspondence with Richard Lenski, 2018-04-27). Ara+ colonies were re-streaked on MA plates, confirmed as Ara+ based on white color when cultured on TA agar medium, and an individual Ara+ colony from each of the five lines was isolated by culturing in DM1000 medium and freeze-storage with glycerol at –80 ºC.

One REL606 colony and its derived Ara+ revertant were confirmed by Sanger sequencing. For this, colony PCR was performed using the following primers for the *ara* operon[31]: REL256 (coordinates for REL606: 70,660–70,783) 5'-CCGATACGCTCATGGGCTTGTTTA-3' and REL257 (coordinates: 71,177–71,154) 5'-CTGCCCAGGCCGTTGCGACTCTAT-3' . To release DNA for the PCR reaction, 2–3 colonies (Ø 2 mm) from plates containing 24–48 h pure cultures were transferred to 50 $\mu$L of 20 mM NaOH, boiled for 10 min at 100 ◦C, spun down, vortexed, and spun down again. PCR was performed with 2 $\mu$L of the resultant solution for use as template DNA, 1.25 U of Taq DNA Polymerase (New England Biolabs, Ipswitch, MA, USA), 0.2 $\mu$M each of the *ara* primers, and 200 $\mu$M of dNTPs in a final volume of 50 $\mu$L of 1 × Standard Taq Reaction Buffer. The cycling conditions were as follows: 95 ºC for 5 min to enhance cell lysis, and 30 cycles of 95 ºC for 30 s, 56 ºC for 30 s and 72 ºC for 1 min, with a final extension at 72 ºC for

5 min. The PCR products were Sanger sequenced by a third party (Institute of Biotechnology, University of Helsinki, Finland) using in-house protocols. The sequences were assembled from Sanger sequencing chromatograms using Pregap4 and Gap4 in the Staden Package[32], and aligned with ClustalW using MEGA7[33] (Fig. S2).

**Competition assay for ancestral strains: *E. coli* REL606 and REL607-like Ara$^+$ reversion mutant**

To ascertain that there was no initial difference in fitness between the two ancestral strains, a competition assay was carried out following a previously established protocol (barricklab.org/twiki/bin/view/Lab/ProceduresLongTermCompetitions, accessed 2018-05-16). For the assay, the strains were revived by culturing 2 $\mu$L of freeze-stored clonal culture in 5 mL LB liquid medium overnight at 37 °C with constant rotation at 120 r.p.m. Subsequent culturing steps were performed in DM1000 medium. Before starting the competition assay, the strains were mixed together at 200-fold dilutions (100-fold overall dilution in cell number), and this mix was immediately dilution plated on TA agar to determine the initial frequencies of the strains. Subsequently, 50 $\mu$L of each of the strains was added to 10 mL medium, with 12 replicates, followed by culturing at 37 °C, and serially diluting (1 % volume) each day for a total of three days. This allows relative fitness to be determined with a precision of ± 1 % (95 % confidence intervals). Upon completion of the assay, the cultures were immediately dilution plated on TA agar to determine the final frequencies of the strains.

The relative fitness ($W$) of the strains was calculated as the ratio of their Malthusian parameters ($M_{REL606}$ and $M_{REL607\text{-}like}$): $W = \frac{\log(M_{REL606})}{\log(M_{REL607-like})}$. The Malthusian parameters are given by $M_{REL606} = \frac{N_{REL606}(f)}{N_{REL606}(i)} = \frac{PC_{REL606}(f) \cdot DF}{PC_{REL606}(i)}$ and $M_{REL607-like} = \frac{N_{REL607-like}(f)}{N_{REL607-like}(i)} = \frac{PC_{REL607-like}(f) \cdot DF}{PC_{REL607-like}(i)}$, where $N$ is the cell number, $PC$ the plate count on TA agar, $DF$ the dilution factor of all transfers combined, and $i$ and $f$ the initial and final time points, respectively. Therefore, a difference between the logarithm of the Malthusian parameters (ratio ≠ 1) for the two strains (paired for each replicate) indicates a difference in fitness. The result of the competition assay (one of 12 replicates was lost due to technical error, *i.e.* $N$ = 11) supports the absence of a difference in fitness between the two ancestral strains (Fig. S3).

Fig. S3 was created using the method by[34] (tool available at http://www.estimationstats.com, accessed 2019-07-19). The paired mean difference between log($M_{REL606}$) and log($M_{REL607\text{–}like}$) is -0.0056 [95.0 % CI -0.0488, 0.0418]. The two-sided *P*-value of the Wilcoxon test is 0.594. The effect sizes and CIs are reported above as: effect size [*CI width lower bound; upper bound*]. 5000 bootstrap samples were taken; the confidence interval is bias-corrected and accelerated. The *P*-value reported is the likelihood of observing the effect size if the null hypothesis of zero difference is true.

**Details on serial passage experiment**

The experiment was started by culturing the REL606 and REL607-like strains in DM1000 for 24 h to obtain 2 × $10^9$ cells $mL^{-1}$. One colony from each plated strain was added to 2 × 5 mL of DM1000, and from this solution, approximately $10^6$ bacterial cells (50 $\mu$L) of the corresponding strain were pipetted to the wells of the deep well plates containing 500 $\mu$L of DM1000 with 0.5 × MIC of the appropriate antimicrobial. In addition, 5 × $10^5$ plaque forming units (PFU) of the T4 phage (13 $\mu$L from phage stock containing 3.9 × $10^9$ PFU $mL^{-1}$) were pipetted to phage treatment wells. Deep well plates were cultured for at 37 °C with constant shaking at approx. 120 r.p.m. Serial passage was performed every 24 h by transferring 10 $\mu$L (2 % v/v) from each well to a new well containing fresh medium using a multichannel pipette. To store populations in suspended animation, after the first 48 h and subsequently every 96 h in the experiment, all deep well plates were freeze-stored at –20 °C by mixing in 250 $\mu$L of sterile 85 % glycerol in each well and covering with a foil seal.

**Determining minimum inhibitory concentrations for antimicrobials for ancestral *E. coli* strain**

The MIC values for the experimental antimicrobials were determined for the REL606 strain in the experimental conditions using the microdilution method. Instead of 2-fold dilutions typically used, the experiment was performed at even concentration intervals in a total interval determined based on expected MIC values: for nalidixic acid, 1–5 $\mu$g $mL^{-1}$ (expected MIC of $\mu$g $mL^{-1}$ for REL606 based on[35]; for rifampicin, 1–5 $\mu$g $mL^{-1}$ (expected MIC of 2.5 $\mu$g $mL^{-1}$ for *E. coli* B[36]); for spectinomycin, 10–20 $\mu$g $mL^{-1}$ (expected value of 10–20 μg mL$^{-1}$ for *E. coli* B[37]). The experiment was performed by adding $10^6$ cells (1/2000 dilution of overnight culture of REL606 in DM1000 medium) to honeycomb wells containing DM1000 medium, with each well containing a different concentration of the antimicrobial in question according to the gradients specified above. This cell number is expected to be lower than the spontaneous frequency of resistance mutations, indicating that any observed growth represents the intrinsic resistance level of the ancestral strain rather than that of derived mutants emerging during the test. Culturing was performed for 24 h at 37 °C using the Bioscreen C well-plate reader (Labsystems, Helsinki, Finland) that measures optical density (OD) at 420–580 nm with a wideband filter at 5 min intervals.

The MIC value was interpreted as the smallest concentration preventing growth, which was 1.0, 2.6 and 8.0 μg mL$^{-1}$ (experimental concentrations of 0.5 × MIC thereby being 0.5, 1.3 and 4.0 μg mL$^{-1}$) for nalidixic acid, rifampicin, and spectinomycin, respectively (Fig. S7). The dose response curves in Fig. S7 show that the antimicrobial level where the carrying capacity begins to decrease is lower than the concentration (0.5 × MIC) used in the experiment. This indicates that differences in effective population size caused by the antimicrobials do not account for differences in evolutionary dynamics between the treatments in the absence of evolution altering the population size response to the antimicrobials.

**Details on phenotyping experimental populations over time**

To track the development of antimicrobial resistance over time, the following selective concentrations were chosen for each antimicrobial for use in LB agar plates based on literature: 2 and 20 μg mL$^{-1}$ for nalidixic acid, 50 μg mL$^{-1}$ for rifampicin, and 50 and 75 μg mL$^{-1}$ for spectinomycin. After each 24 h growth cycle in the experiment, the populations were transferred to selective plates, as well as plates without antimicrobials to control for the presence of bacteria, by pin-replicating the whole deep well plate onto a large petri dish. To assess the presence of phage, 200 μL of overnight culture of the ancestral REL606 strain into was added to a sterile 15 mL Eppendorf tube, followed by adding 15 mL of soft agar tempered to 55 °C in a water bath, vortexing, pouring on large petri dish containing LB agar, and pin-replication of the 24 h culture from the experiment upon solidification of soft agar. Plates were cultured at for 24 h at 37 °C. Each plate was documented by photography, and the results were interpreted as negative (0), weak

(1), or strong (2) bacterial growth (antimicrobial resistance) or plaque formation (phage presence), subsequently converted to binary data (2 converted to 1).

**Details on phenotyping clones isolated from experimental end-point**

Clones were isolated from the experimental end-point by streaking an inoculum from surviving populations ($N$ = 900 out of 928 populations in total) on LB plates. Following culture (24 h / 37 °C), a single colony was selected and transferred to a 96 well plate containing 200 $\mu$L of DM1000. Following culture (24 h / 37 °C / 120 r.p.m.), the clonal lines were freeze-stored with 85 % glycerol at –80 °C. To perform subsequent tests, the master well plates containing the original clones were replicated on 96 well plates containing 200 $\mu$L of DM1000. The test well plates were cultured and freeze-stored as above. To obtain quantitative growth traits (optical density at 600 nm) for different antimicrobial concentrations, a test well plate was replicated on another 96 well plate containing 200 $\mu$L of DM1000, and cultured for 24 h at 37 °C / 120 r.p.m. This results in a cell density of 2 × $10^9$ cells $mL^{-1}$. The clonal cultures were diluted into deep well plates to obtain a total dilution of 1:2000, corresponding to 1 × $10^6$ cells $mL^{-1}$ assuming a density of 2 × $10^9$ cells $mL^{-1}$. This cell count is expected to be lower than the frequency of spontaneous antimicrobial resistance mutations, indicating that any observed growth represents the (potentially evolved) resistance level of the clonal line rather than that of derived mutants emerging during the test. Approx. 2000 cells (2 $\mu$L) of each diluted clone was pipetted on another 96 well plate containing 300 $\mu$L of DM1000 with each of the experimental antimicrobials at the following concentrations: 0, 1, 2, 10, or 100 × MIC (respectively: 0, 1.0, 2.0, 10, and 100 $\mu$g $mL^{-1}$ for nalidixic acid; 0, 2.6, 5.2, 26, and 260 $\mu$g $mL^{-1}$ for rifampicin; and 0, 8.0, 14, 80, and 800 $\mu$g $mL^{-1}$ for spectinomycin. Following culture for 24 h at 37 °C / 120 r.p.m., the cell density reached by each clone was quantified by measuring OD at 600 nm wavelength (Tecan Infinite M200) with a bandwidth of 9 nm and 25 measurements per well. The REL606 and REL607-like strains and blank wells containing the medium alone supplemented with the different antimicrobials (only rifampicin was shown to alter the background OD level) were used as controls in each well plate. To determine the phage resistance phenotype of the clones from the phage treatments, a work plate containing REL606 and REL607-like strains as controls was pin-replicated on an LB plate containing a top layer of soft agar with the ancestral T4 phage (see detailed protocol for plaque assay above). Following culture for 24 h at 37 °C, phage resistance was quantified for each clone as a binary trait based on the absence (0) or presence (1) of bacterial growth on the phage lawn.

**Details on extraction of DNA from clones isolated from experimental end-point**

DNA extraction was performed from 1 mL of overnight culture with the Qiaqen DNeasy 96 Blood & Tissue kit using a custom protocol. To release nucleic acids, the cells were transferred on S plates and spun down for 30 min at 6200 r.p.m., followed by addition of 200 $\mu$L of ATL and proteinase K solution (prepared by mixing 36 mL of ATL and 4 mL of proteinase K) to each well, vortexing for 15 s with plastic cover attached, and briefly spinning down at 3000 r.p.m. The suspension was incubated for 30 min at 56 °C, with vortexing every 10 min. Subsequently, 410 $\mu$L of AL solution was added to each well, followed by vortexing for 15 s, and spin down at 3000 r.p.m. as described above. The whole sample (approx. 620 $\mu$L) was transferred to the extraction membrane columns, followed by spinning down for 10 min at 6000 r.p.m. in room temperature (RT) with plastic cover attached and discarding the flow-through. Subsequently, 500 $\mu$L of AW1 was added, followed by spinning down at 6000 r.p.m. in RT and discarding the flow-through. The plastic cover was removed, followed by adding 500 $\mu$L of AW2, spinning down for 15 min at 6000 r.p.m. in RT, discarding the flow-through, and centrifuging empty wells for 2 min at 6000 r.p.m. in RT to remove ethanol residues. After the ethanol was allowed to evaporate for 2 min in RT, membrane columns were transferred to elution plates. Elution was performed in DNA-free grade water by incubating in 40 $\mu$L of elute for 2 min in RT, centrifuging for 2 min at 6000 r.p.m., adding another 40 $\mu$L of sterile DNA-free grade water and centrifuge immediately for 2 min at 6000 r.p.m. The extracted DNA was stored at –20 °C.

### Supplementary results

##### Details on potential molecular targets of resistance

*Nalidixic acid resistance*: Nalidixic acid exposure was associated with a small number of nonsynonymous mutations overall (median of 0 mutations in time-invariant single agent environment in the absence of phage, equivalent to control environment) as well as a relatively small number of recurrent mutational targets occurring only in a small proportion of the isolates (Fig. 2G). This is consistent with major clones lacking resistance in time-invariant nalidixic acid environment in the absence of phage (Fig. 2C). These targets include *acrR* (encoding HTH-type transcriptional regulator), *rfaQ* (lipopolysaccharide core heptosyltransferase) and ECB RS03400 (putative phosphoglucomutase). Among these genes, *acrR* has been previously implicated in quinolone resistance[38], while the product function (LPS biosynthesis) and previous findings suggest that *rfaQ*[39] and phosphoglucomutase[40,41] may be associated with resistance to both quinolone and phage.

*Rifampicin resistance*: The vast majority of clones from the time-invariant single agent environment had nonsynonymous mutations (median of 1.5 mutations in the absence of phage) in the gene *rpoB* (β subunit of RNA polymerase) known to produce rifampicin resistance[42], and the presence of mutations in this gene was almost exclusively associated with a resistance phenotype (Fig. 2G). In addition, five other genes (*galU*, *infB*, *marR*, *mreC* and *mrdB*) were recurrently mutated only in the presence of rifampicin. Among these, *marR* is related to drug efflux[43]; *mreC* and *mrdB* are both related to cell shape, with *mreC* mutations having been previously associated with rifampicin exposure[44]; and mutations in the essential gene *infB* (translation initiation factor 2) have been previously implicated in rifampicin resistance[45] and compensating for the fitness cost of antimicrobial resistance mutations[46]. The gene *galU* is involved in LPS biosynthesis and has been previously implicated in phage resistance[47], and consistent with this, was only hit in the presence of phage in this study.

*Spectinomycin resistance*: The time-invariant spectinomycin (aminoglycoside class) environment, either in the presence or absence of phage, led to higher mean spectinomycin resistance prevalence among end-point clones compared to the antimicrobial-free control and time-invariant combination environments, although resistance increased markedly only in the time-dependent environment in the absence of phage (Fig. 2E; antimicrobial regime, $P < 0.001$; for full results, see Table S5). We were unable to obtain robust population-level time series data for spectinomycin, as the data was flooded in (majority of populations gave) positive signal masking any potential treatment effect. There are a number of factors that could enable population growth in our selective conditions, resulting in a positive resistance signal despite all or most of the cells remaining sensitive. Resistance may occur only in a subset of the population owing to a low selection coefficient, reversible amplifications[48,49], or plastic gene regulatory changes induced by antimicrobial stress (*e.g.* SOS or stringent response)[50,51]. Three genes, *nadR* (transcriptional regulator), *trkH* (potassium uptake protein) and ECB RS09520 (putative carboxyl-terminal processing protease), were recurrently mutated specifically in the presence of spectinomycin (median of 1 nonsynonymous mutation in time-invariant single agent environment), with only *trkH* reaching high (close to 0.5) frequencies (Fig. 2G). Among these, *trkH* has been previously implicated in aminoglycoside resistance[52,53], while *nadR*[54] and proteases[55] can, among other functions, be related to the bacterial stress response. Together these findings suggest that the time-invariant spectinomycin environment may cause weak selection pressure for resistance. As much higher resistance levels occur in a subset of the time-dependent environments, with a shorter spectinomycin selective window, these are expected to arise from the other agents (see results section"Pleiotropic and fitness effects of strong selective agents, rifampicin and phage, modulate evolutionary outcome" in main text).

*Phage resistance*: Phage exposure caused recurrent mutations in several genomic targets almost exclusively associated with phage resistant phenotypes, with most targets exhibiting low to moderate (max. 0.5) prevalence among isolates (Fig. 2G). This indicates that resistance to the phage T4 has a wide target in *E. coli* B. This is consistent with earlier studies showing that membrane modifications preventing phage adsorption represent a highly common resistance mechanism of bacteria against virulent phages and can typically be achieved by mutations in a number of genes affecting membrane structure and components[56]. Moreover, the phage caused an increase of one nonsynonymous mutation in the median mutation count of the clones, suggesting that individual mutations rather than several mutations in combination were required for phage resistance. The potential phage resistance targets discovered in this study encompass 14 genes:

*acrR*, *asmA*, *fabR*, *galU*, *infB*, *lpcA*, *mscM*, *rfaQ*, ECB RS0200, ECB RS03400, ECB RS03925, ECB RS05770, ECB RS09520, and ECB RS18465 (Fig. 2G). Six among these (*acrR*, *galU*, *infB*, *rfaQ*, ECB RS03400 and ECB RS09520) were also associated with resistance to a particular antimicrobial compound and have been discussed above. Among the eight remaining genes specifically associated with phage exposure and phage resistance phenotypes, mutations in four genes (*asmA*, *fabR*, *mscM* and ECB RS18465) were particularly prevalent, with mutations in *mscM* being prevalent across antimicrobial compound environments but absent from the antimicrobial-free control environment. The product of *asmA* is involved in the assembly of outer membrane proteins and has been implicated in phage resistance[57]. The product of *fabR* is an HTH-type transcriptional repressor involved in unsaturated fatty acid biosynthesis and the physical properties of the cell membrane. Recently, both *fabR*[58] and phage resistance[59] have more specifically been linked to L-threonine production. The gene *mscM* encodes a mechanosensitive channel protein located in the cell membrane. The channel exhibits long-term open states in response to increased membrane tension[60], suggesting that loss-of-function mutations may be beneficial in the presence of both phage and antimicrobial compounds if low antimicrobial concentrations increase membrane tension and open states facilitate phage adsorption. Finally, ECB RS18465 encodes a putative glucosyltransferase. Glycosyltransferases have been implicated in bacterial phage resistance through determining the glycosylation pattern of cell membrane teichoic acids required for phage adsorption[61].

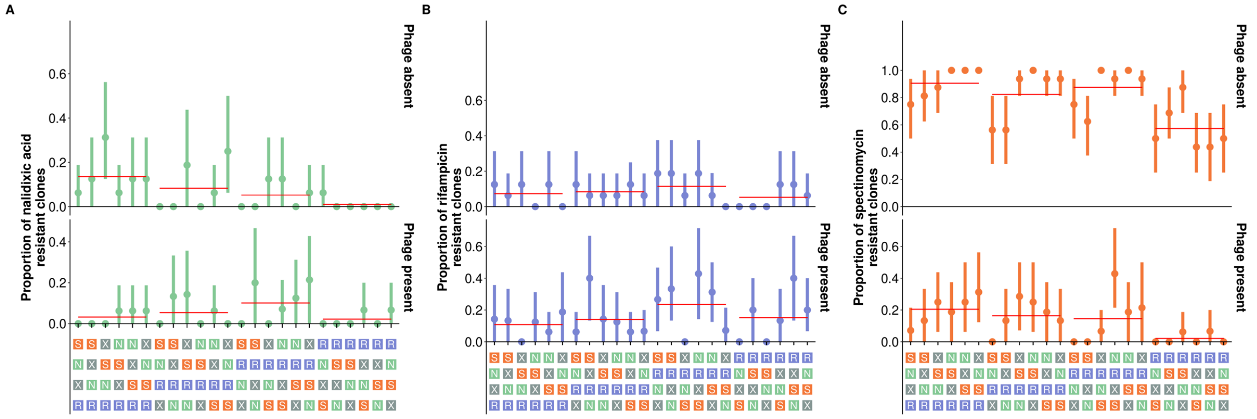


**Supplementary Figure S1. Antimicrobial resistance spectra in different time-dependent regimes. Related to Figure 2, Figure 3, Figure 4, and Figure 5.** (A), (B) and (C) show resistance spectra for nalidixic acid, rifampicin and spectinomycin, respectively, at the experimental end-point depending on antimicrobial sequence and presence of phage (mean resistance ± bootstrapped 95 % confidence intervals). The data is based on *N* = 900 clones isolated from populations at the experimental end point. The antimicrobial sequences have been ordered by the rifampicin exposure epoch explaining resistance to all three antimicrobials, with the mean resistance level indicated by a red line.

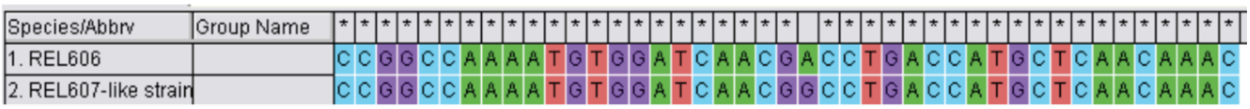


**Supplementary Figure S2. Confirmation of REL607-like Ara+ reversion mutation. Related to Figure 2, Figure 3, Figure 4, and Figure 5.** The figure is based on ClustalW alignment in MEGA7.

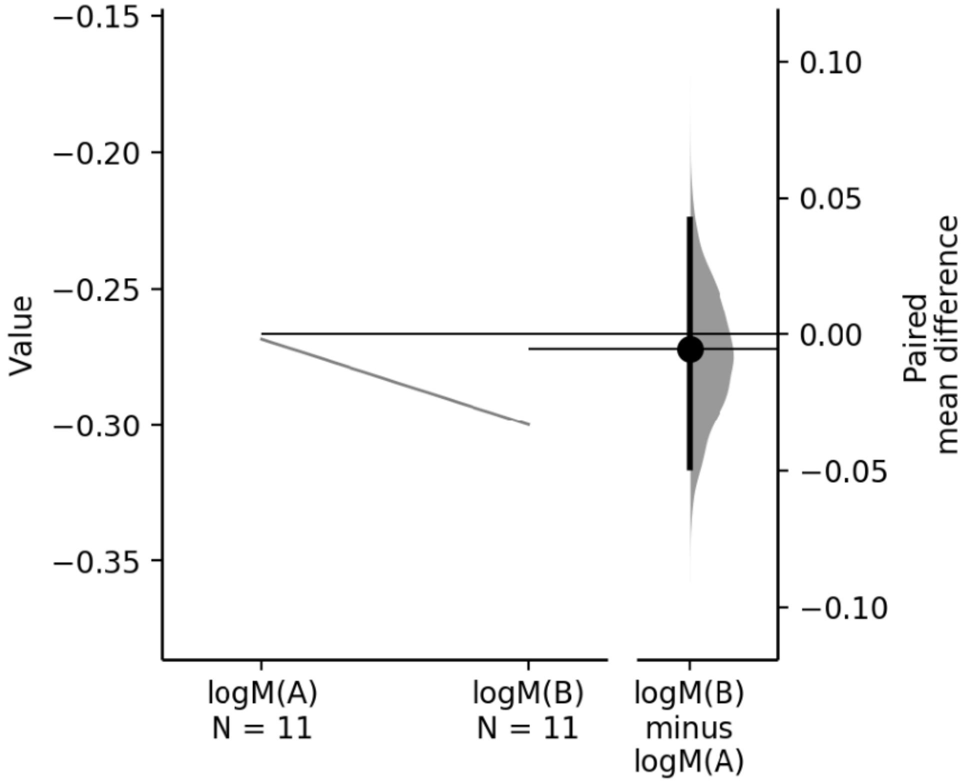


**Supplementary Figure S3. Malthusian parameters for the two ancestral strains in the serial passage experiment showing lack of difference in initial fitness. Related to Figure 2, Figure 3, Figure 4, and Figure 5.** The paired mean difference between log(MREL606) (A) and log(MREL607-like) (B) is shown in the above Gardner-Altman estimation plot. Both groups are plotted on the left axes as a slopegraph: each paired set of observations is connected by a line. The paired mean difference is plotted on a floating axis on the right as a bootstrap sampling distribution. The mean difference is depicted as a dot; the 95 % confidence interval is indicated by the ends of the vertical error bar.

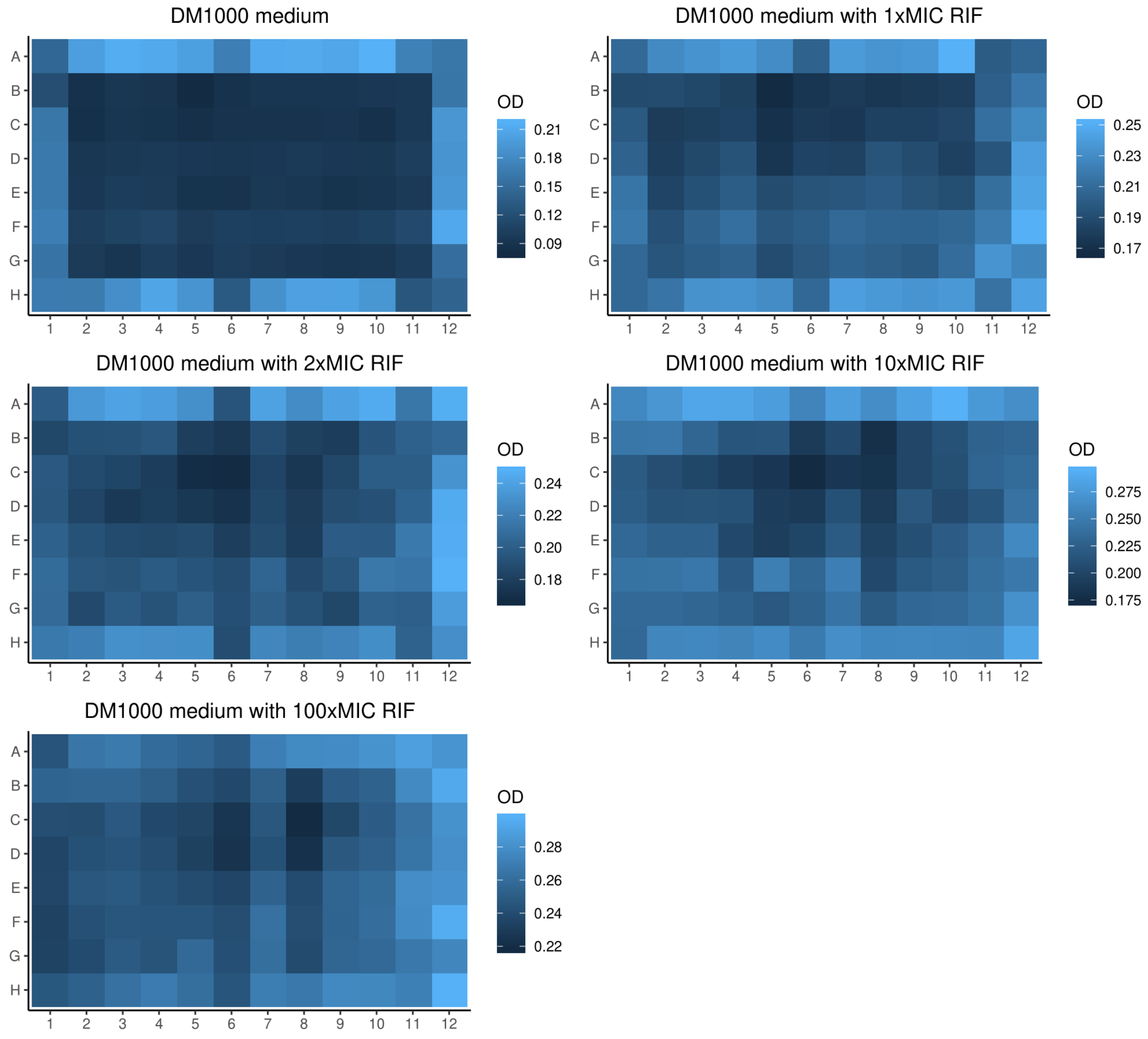


**Supplementary Figure S4. Effect of well plate location on background optical density (OD) value at 600 nm obtained from the Tecan Infinite M200 device used to quantify the growth of end-point clones from serial passage experiment. Related to Figure 2, Figure 3, Figure 4, and Figure 5.** The effect is shown separately for the DM1000 medium alone and with different levels of rifampicin, which affected the background OD level. The heat map displays the mean of 10 measurements per well per condition.

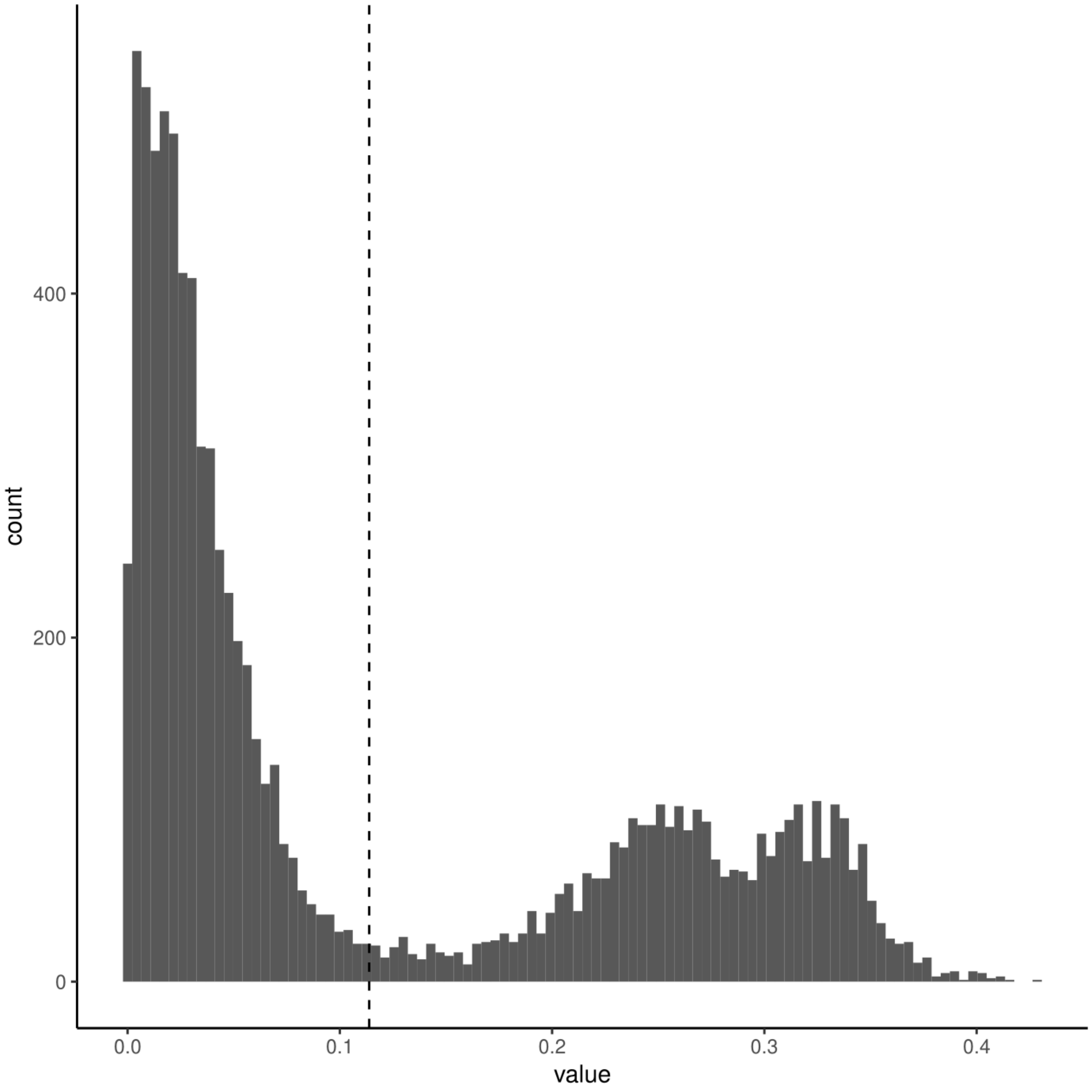


**Supplementary Figure S5. Distribution of OD data for clonal lines after removing well-specific background OD values. Related to Figure 2, Figure 3, Figure 4, and Figure 5.** The *x*-axis displays OD at 600 nm. The cut-off used to remove false positives (3 × standard deviation of lowest distribution mirrored on each side of zero) is indicated with a dashed line. All growth data below the cut-off value was transformed to zero growth.

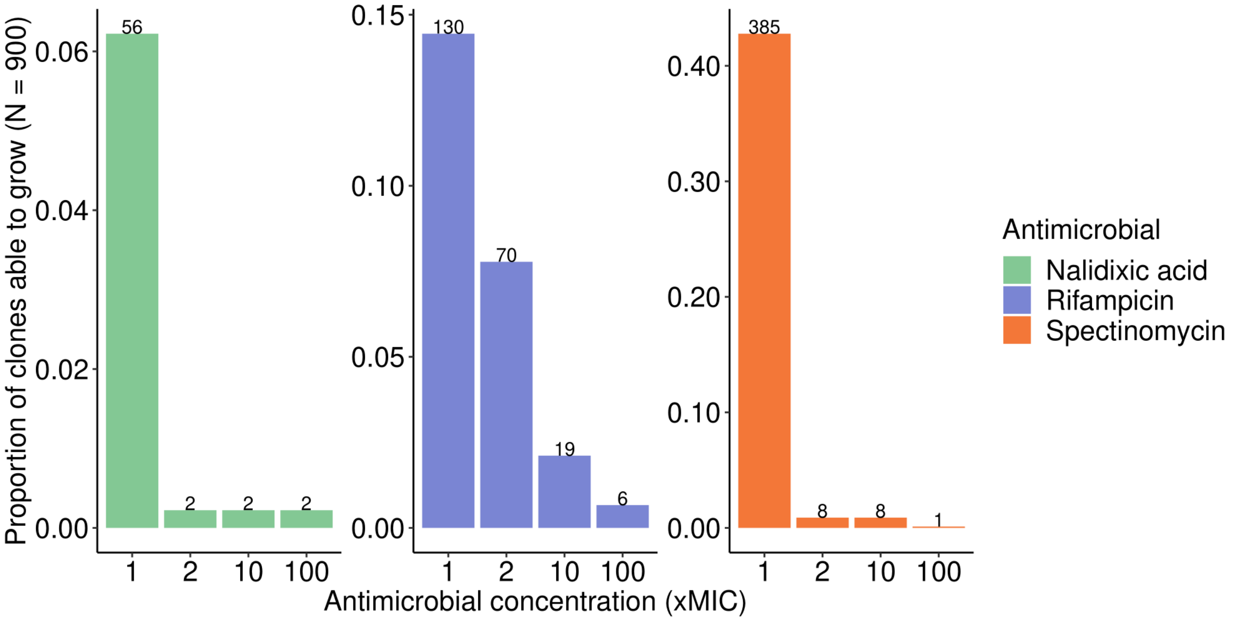


**Supplementary Figure S6. Ability of clones isolated from the end-point of evolutionary experiment to grow at different concentrations of the experimental antimicrobials. Related to Figure 2, Figure 3, Figure 4, and Figure 5.** The *y*-axis shows the proportion of clones and the numbers on top of bars the number of clones ($N = 900$).

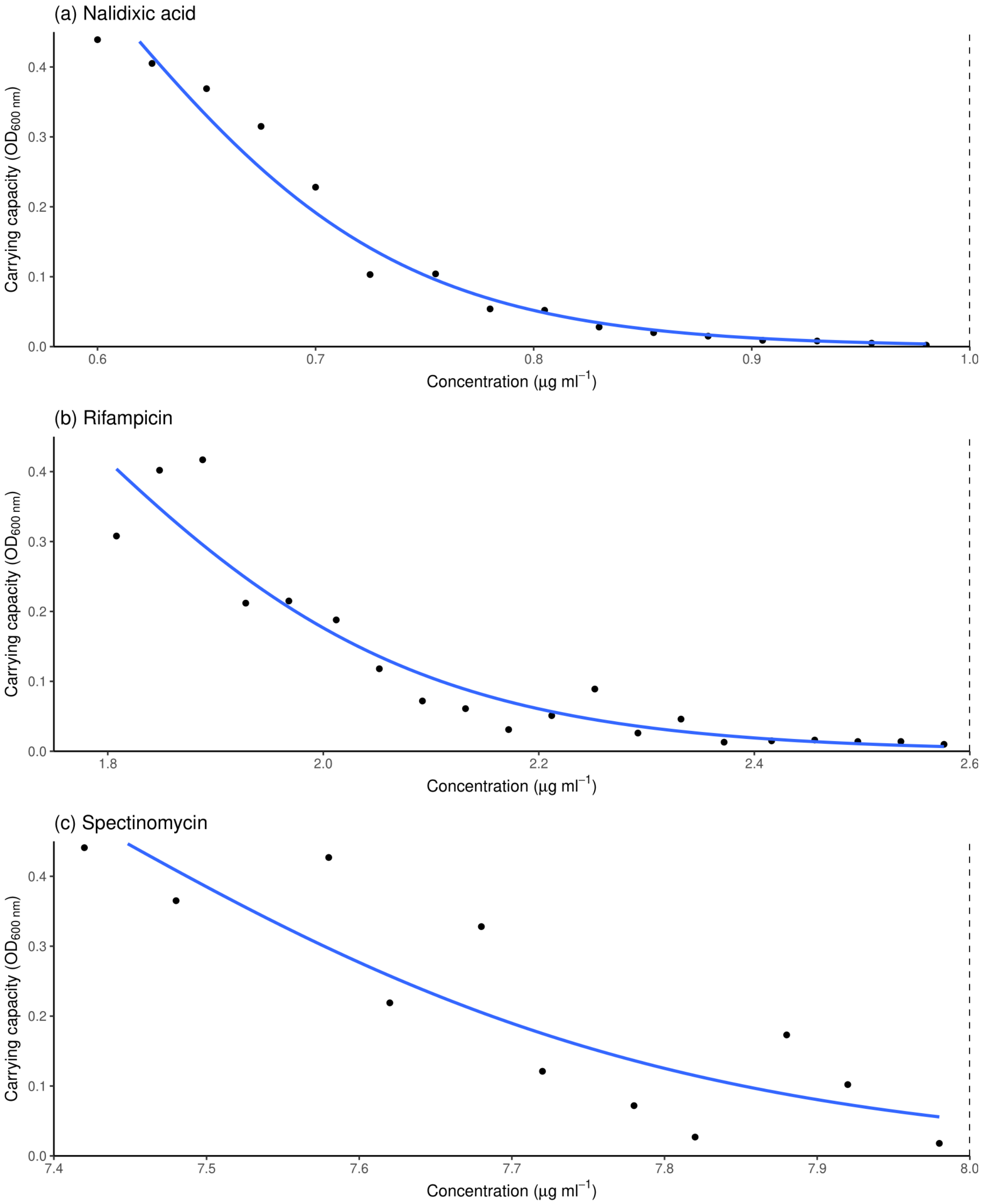


**Supplementary Figure S7. Dose response curve for REL606 carrying capacity in the experimental antimicrobials. Related to Figure 2, Figure 3, Figure 4, and Figure 5.** (A) nalidixic acid, (B) rifampicin, and (C) spectinomycin. The minimum inhibitory concentration (MIC) where growth is no longer observed is shown with a dashed line. The blue line is a logistic regression curve fitted to the data.

**Supplementary Table S1. Related to Figure 2, Figure 3, Figure 4, and Figure 5.** ANOVA table for generalized least squares model on temporal nalidixic acid resistance dynamics at population level.

| Model term | *Df* | *F* | *P* |
|---:|---:|---:|---:|
| Antimicrobial regime (control / time-invariant single-agent / time-invariant combination / time-dependent) | 3 | 29.3 | < 0.001 |
| Phage (present/absent) | 1 | 32.4 | < 0.001 |
| Antimicrobial regime × phage | 3 | 0.39 | 0.76 |
| Residuals | 3348 | | |

**Supplementary Table S2. Related to Figure 2, Figure 3, Figure 4, and Figure 5.** ANOVA table for binomial generalized linear (*i.e.* logistic regression) model on nalidixic acid resistance levels in end-point clones.

| Model term | *Df* | Deviance | Residual *df* | $P(\chi^2)$ |
|---:|---:|---:|---:|---:|
| Antimicrobial regime (control / time-invariant single-agent / time-invariant combination / time-dependent) | 3 | 11.9 | 835 | 0.008 |
| Phage (present/absent) | 1 | 0.041 | 834 | 0.84 |
| Antimicrobial regime × phage | 3 | 6.80 | 831 | 0.079 |

**Supplementary Table S3. Related to Figure 2, Figure 3, Figure 4, and Figure 5.** ANOVA table for generalized least squares model on temporal rifampicin resistance dynamics at population level.

| **Model term** | ***Df*** | ***F*** | ***P*** |
|---:|---:|---:|---:|
| Antimicrobial regime (control / time-invariant single-agent / time-invariant combination / time-dependent) | 3 | 29.3 | < 0.001 |
| Phage (present/absent) | 1 | 5.49 | 0.019 |
| Antimicrobial regime × phage | 3 | 1.58 | 0.19 |
| Residuals | 3340 | | |

**Supplementary Table S4. Related to Figure 2, Figure 3, Figure 4, and Figure 5.** ANOVA table for binomial generalized linear (*i.e.* logistic regression) model on rifampicin resistance levels in end-point clones.

| Model term | *Df* | Deviance | Residual *df* | $P(\chi^2)$ |
|---:|---:|---:|---:|---:|
| Antimicrobial regime (control / time-invariant single-agent / time-invariant combination / time-dependent) | 3 | 114.1 | 833 | < 0.001 |
| Phage (present/absent) | 1 | 13.5 | 832 | < 0.001 |
| Antimicrobial regime × phage | 3 | 0.61 | 829 | 0.89 |

**Supplementary Table S5. Related to Figure 2, Figure 3, Figure 4, and Figure 5.** ANOVA table for binomial generalized linear (*i.e.* logistic regression) model on spectinomycin resistance levels in end-point clones.

| **Model term** | ***Df*** | **Deviance** | **Residual *df*** | $P(\chi^2)$ |
|---|---|---|---|---|
| Antimicrobial regime (control / time-invariant single-agent / time-invariant combination / time-dependent) | 3 | 35.6 | 836 | < 0.001 |
| Phage (present/absent) | 1 | 327.3 | 835 | < 0.001 |
| Antimicrobial regime × phage | 3 | 32.1 | 832 | < 0.001 |

**Supplementary Table S6. Related to Figure 2, Figure 3, Figure 4, and Figure 5.** ANOVA table for binomial generalized linear (*i.e.* logistic regression) model on phage resistance levels in end-point clones.

| Model term | *Df* | Deviance | Residual *df* | *P* ($\chi^2$) |
|---|---|---|---|---|
| Antimicrobial regime (control / time-invariant single-agent / time-invariant combination / time-dependent) | 3 | 17.9 | 440 | $< 0.001$ |

**Supplementary Table S7. Related to Figure 2, Figure 3, Figure 4, and Figure 5.** ANOVA table for binomial generalized linear (*i.e.* logistic regression) model on spectinomycin resistance levels in end-point clones comparing control and time-invariant rifampicin environments.

| Model term | *Df* | Deviance | Residual *df* | $P(\chi^2)$ |
|---|---|---|---|---|
| Antimicrobial regime (control / time-invariant rifampicin environment) | 1 | 4.24 | 59 | 0.039 |
| Phage (present/absent) | 1 | 13.3 | 58 | $<0.001$ |
| Antimicrobial regime × phage | 1 | 9.97 | 57 | 0.002 |

**Supplementary Table S8. Related to Figure 2, Figure 3, Figure 4, and Figure 5.** ANOVA table for binomial generalized linear (*i.e.* logistic regression) model on rifampicin resistance levels in end-point clones as a function of rifampicin exposure epoch and phage exposure.

| Model term | *Df* | Deviance | Residual *df* | $P(\chi^2)$ |
|---|---|---|---|---|
| Rifampicin exposure epoch | 3 | 6.94 | 746 | 0.074 |
| Phage (present/absent) | 1 | 11.2 | 745 | < 0.001 |
| Rifampicin exposure epoch × phage | 3 | 1.24 | 742 | 0.74 |

**Supplementary Table S9. Related to Figure 2, Figure 3, Figure 4, and Figure 5.** ANOVA table for binomial generalized linear (*i.e.* logistic regression) model on nalidixic acid resistance levels in end-point clones as a function of nalidixic acid and rifampicin exposure epochs in the absence of phage exposure.

| Model term | *Df* | Deviance | Residual *df* | $P(\chi^2)$ |
|---|---|---|---|---|
| Nalidixic acid exposure epoch | 3 | 2.46 | 380 | 0.48 |
| Rifampicin exposure epoch | 3 | 13.9 | 377 | 0.003 |
| Nalidixic acid exposure epoch × rifampicin exposure epoch | 5 | 4.93 | 372 | 0.42 |

**Supplementary Table S10. Related to Figure 2, Figure 3, Figure 4, and Figure 5.** ANOVA table for binomial generalized linear (*i.e.* logistic regression) model on nalidixic acid resistance levels in end-point clones as a function of nalidixic acid and rifampicin exposure epochs in the presence of phage exposure.

| Model term | *Df* | Deviance | Residual *df* | $P(\chi^2)$ |
|---:|---:|---:|---:|---:|
| Nalidixic acid exposure epoch | 3 | 12.2 | 362 | 0.007 |
| Rifampicin exposure epoch | 3 | 4.37 | 359 | 0.22 |
| Nalidixic acid exposure epoch × rifampicin exposure epoch | 5 | 10.2 | 354 | 0.069 |

**Supplementary Table S11. Related to Figure 2, Figure 3, Figure 4, and Figure 5.** ANOVA table for binomial generalized linear (*i.e.* logistic regression) model on spectinomycin resistance levels in end-point clones as a function of spectinomycin and rifampicin exposure epochs and phage exposure.

| Model term | *Df* | Deviance | Residual *df* | $P(\chi^2)$ |
|---:|---:|---:|---:|---:|
| Spectinomycin exposure epoch | 3 | 11.0 | 746 | 0.012 |
| Rifampicin exposure epoch | 3 | 47.5 | 743 | $<0.001$ |
| Phage (presence/absence) | 1 | 416.0 | 742 | $<0.001$ |
| Spectinomycin exposure epoch × rifampicin exposure epoch | 5 | 10.2 | 737 | 0.071 |
| Spectinomycin exposure epoch × phage | 3 | 1.74 | 734 | 0.63 |
| Rifampicin exposure epoch × phage | 3 | 1.41 | 731 | 0.70 |
| Spectinomycin exposure epoch × rifampicin exposure epoch × phage | 5 | 5.96 | 726 | 0.31 |

**Supplementary Table S12. Related to Figure 2, Figure 3, Figure 4, and Figure 5.** Davis-Mingioli medium with 1000 mg $L^{-1}$ glucose (DM1000) for culturing strains in liquid medium during the competition assay, serial passage experiment, and clone measurements. Culture conditions: 24 h at 37 °C. The medium is prepared by adding $dH_2O$ to final volume and autoclaving, followed by adding 0.5 mL each of the following stock solutions: 10 % (w/v) magnesium sulfate $MgSO_4$ (separately autoclaved stock) and 0.2 % (w/v) thiamine (vitamin B1; filter sterilized), and 5 mL of 10 % glucose (separately autoclaved stock). Sources: Levin, et al. [62]; Lenski lab website (myxo.css.msu.edu/ecoli/dm25liquid.html, accessed 2018-05-04); and Barrick lab website (barricklab.org/twiki/bin/view/Lab/ProtocolsRecipesDavisMingioli, accessed 2018-06-20; barricklab.org/twiki/bin/view/Lab/ProtocolsFluctuationTests, accessed 2018-06-26).

| Component | 0.5 L |
|---|---|
| Potassium phosphate (dibasic) $K_2HPO_4$* | 2.65 g |
| Potassium phosphate (monobasic anhydrous) $KH_2PO_4$ | 1 g |
| Ammonium sulfate $(NH_4)_2SO_4$ | 0.5 g |
| Sodium citrate (trisodium, dihydrate) $Na_3C_6H_5O_7 \times 2(H_2O)$ | 0.25 g |

*If using potassium phosphate (dibasic) trihydrate ($K_2HPO_4 \times 3H_2O$), use 7 g $L^{-1}$.

**Supplementary Table S13. Related to Figure 2, Figure 3, Figure 4, and Figure 5.** Defined minimal arabinose (MA) agar for obtaining new $Ara^{+}$ mutant from $Ara^{-}$ parent. Culture conditions: 48 h at 37 °C. The water is split into three parts, salts, agar, and sugar, which are autoclaved separately and combined, followed by the addition of 1 mL of each of the following stock solutions: 10 % (w/v) magnesium sulfate $MgSO_4$ (separately autoclaved stock) and 0.2 (w/v) thiamine (vitamin B1; filter sterilized). Source: Lenski lab website (http://myxo.css.msu.edu/ecoli/dmagar.html, accessed 2018-05-04).

| **Component** | **1 L** |
|---|---|
| Potassium phosphate (dibasic) $K_2HPO_4$* | 5.3 g |
| Potassium phosphate (monobasic anhydrous) $KH_2PO_4$ | 2 g |
| Ammonium sulfate $(NH_4)_2SO_4$ | 1 g |
| Sodium citrate (trisodium, dihydrate) $Na_3C_6H_5O_7 \times 2(H_2O)$ | 0.5 g |
| $dH_2O$ | 1000 mL |
| Agar | 16 g |
| Antifoam (5 %) | 1 mL |
| l(+)Arabinose | 4 g |

*If using potassium phosphate (dibasic) trihydrate ($K_2HPO_4 \times 3H_2O$), use 7 g $L^{-1}$.

**Supplementary Table S14. Related to Figure 2, Figure 3, Figure 4, and Figure 5.** Recipe for tetrazolium and arabinose (TA) agar plates for distinguishing between Ara$^{-}$ (red/purple) and Ara$^{+}$ (white/pink) strains. Culture conditions: 24 h at 37 °C. The medium is prepared by splitting the water, autoclaving the arabinose separately, combining the two parts after autoclaving, adding TTC from a sterile stock solution, and mixing. Sources: Carlton and Brown [63] and Lenski lab website (http://myxo.css.msu.edu/ecoli/taagar.html, accessed 2018-05-04).

| **Component** | **1 L** | |
|---|---:|---|
| Tryptone | 10 g | |
| Yeast extract | 1 g | |
| Sodium chloride NaCl | 5 g | |
| Agar | 16 g | |
| Antifoam (5 %) | 1 mL | |
| d$H_2O$ | 1000 mL | |
| l(+)Arabinose | 10 g | |
| TTC (5 %) $C_{19}H_{15}ClN_4$ | 1 mL | TTC is the tetrazolium indicator dye |
